# Behavioral algorithms of ontogenetic switching in larval and juvenile zebrafish phototaxis

**DOI:** 10.1101/2025.06.13.659371

**Authors:** Maxim Q. Capelle, Katja Slangewal, Panagiotis E. Eleftheriadis, Armin Bahl

**Affiliations:** Department of Biology, University of Konstanz, 78464 Konstanz, Germany; Centre for the Advanced Study of Collective Behaviour, University of Konstanz, 78464 Konstanz, Germany; Zukunftskolleg, University of Konstanz, 78464 Konstanz, Germany; International Max Planck Research School for Quantitative Behaviour, Ecology and Evolution (IMPRS/QBEE); Max Planck Institute of Animal Behavior, 78315 Radolfzell, Germany

**Keywords:** Zebrafish (*Danio rerio*), behavior, phototaxis, ontogeny, gradient navigation, agent-based simulations

## Abstract

Animals undergo major behavioral adjustments during ontogeny, but how the underlying cognitive algorithms change during this process remains elusive. Here, we describe that zebrafish shift from light-seeking to dark-seeking, as they grow from larval to juvenile stage, within the first few weeks of their life. We apply a combination of complementary phototaxis assays in virtual reality and modeling to dissect the computational basis of this transition. We identify three parallel pathways, one analyzing ambient whole-field luminance levels, one spatially comparing light levels across the eyes, and one computing eye-specific temporal derivatives. Larvae mostly use the latter two spatio-temporal computations for navigation, while juveniles largely employ the first one. We build a library of agent-based models to predict animal behavior across stimulation conditions and in more complex environments. Model-based extraction of latent cognitive variables points towards potential neural correlates of the observed behavioral inversion and illustrates a novel way to explore the processes of vertebrate ontogeny. We suggest that zebrafish phototaxis is regulated via parallel processing streams, which could be a universal implementation to change strategies depending on developmental stage, context, or internal state, making behavior flexible and goal-oriented.

**GRAPHICAL ABSTRACT:** 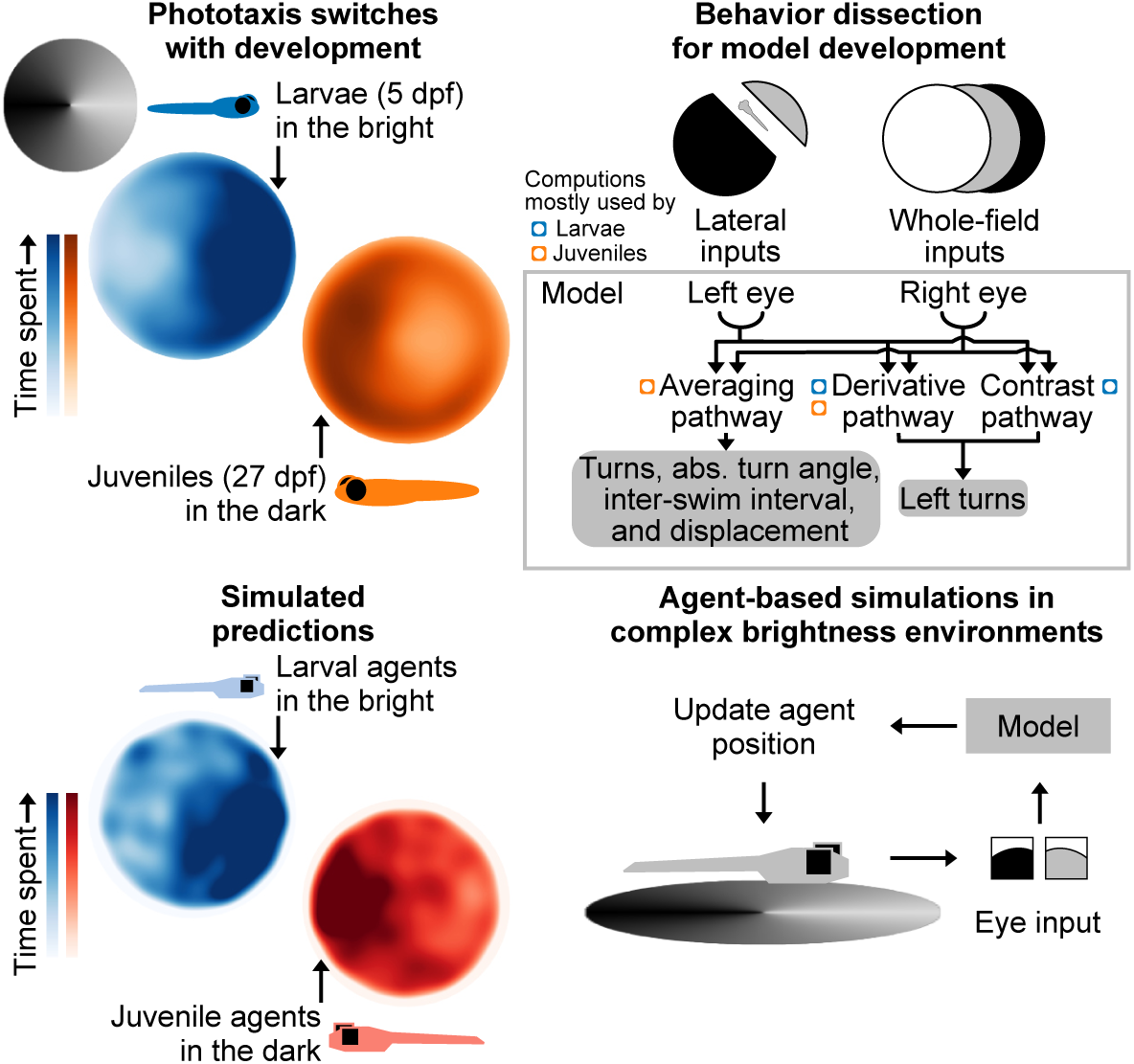

**HIGHLIGHTS:**

- We provide a framework to describe how behavioral strategies evolve during ontogeny.
- Zebrafish switch their phototactic behavior within the first weeks of their life.
- Brightness navigation strategies can be described with a three-pathway model.
- Larvae use spatial cues, while juveniles use ambient whole-field luminance for phototaxis.

## INTRODUCTION

Animal behavior changes during development and in response to different environmental constraints. For example, growing up from infancy to adulthood, humans undergo a long phase of immaturity in which they crucially depend on others for food and protection^1^. Rodent reflexive responses, anxiety behavior, and memory formation change with development^2–4^. Locusts display increased threat-induced hiding behavior over early-life stages^5^. While *Drosophila* larvae avoid light^6,7^, adult *Drosophila* are attracted by light^8,9^. It remains unknown whether such ontogenetic behavioral transitions involve remodeling circuit mechanisms or if they can be explained through adjustments of the algorithmic strategies underlying sensory-motor decision-making.

An excellent system for the study of ontogeny is the vertebrate zebrafish^10^. This freshwater fish shows rapid behavioral adjustments in early development, while continuously interacting with the environment and without the need for parental care. These features provide unique advantages compared to insects and mammalian systems. The majority of insects undergo fundamental structural changes during metamorphosis. While some neural structures remain intact, most of the insect nervous system and body plan get completely remodeled^11^, making it challenging to interpret insect ontogeny. Mammalian development, in general, requires complex social interactions with the parental generation^12^, largely complicating experimentation.

The visual system and motor control elements of zebrafish become fully functional a few days after hatching^13–16^, allowing animals to perform a large range of visually driven behaviors^17^. Around three weeks of age, zebrafish start to engage in social interactions^18–23^ and display robust learning and memory abilities^24–26^. In addition to these behavioral transitions, it has been shown that even certain innately present sensory-motor responses, like the optomotor response and phototaxis, may invert while the animals grow from larval to juvenile to adult stage^27–30^. However, quantitative behavioral analyses of such inversions are largely absent.

In particular, phototaxis—the tendency to alter behavioral responses to light—is well-suited for the study of ontogeny^10^. Luminance stimuli are easy to control, allow for rich variations of spatial and temporal features, and can be used to probe complex gradient tracking strategies and search behavior. Larval zebrafish display positive phototaxis by approaching brighter regions^31,32^. Many studies have explored potential behavioral algorithms and neural implementations of this behavior. When presented with *lateral brightness stimuli*, locked to the position and orientation of the fish, larvae swim towards the brighter side^33,34^, which suggests that spatial contrast across the eyes plays a major role in phototaxis. Temporal luminance dynamics have been shown to provide additional cues for gradient navigation. In response to sudden decreases in whole-field brightness, animals swim more vigorously, and with larger and more frequent turns^32,35–38^. These behaviors can be modulated by adapting animals to different brightness levels^35,39,40^, indicating that phototaxis flexibly depends on the recent environmental context. With advances in virtual reality for high-throughput zebrafish experimentation^21^, it is now becoming possible to extend behavioral assays towards more complex stimulus arrangements and beyond the larval stage.

Quantitative behavioral experiments and computational modeling allow for the extraction and validation of the algorithmic processes underlying ontogeny. In addition, they assist in the mapping of how corresponding cognitive parameters change over time. Such analyses provide a theoretical basis to interpret potential ecologically relevant reasons for the observed changes and guide experiments to assess their mechanistic implementation^41,42^. Several models have been suggested to explain phototaxis in larval zebrafish: Fish may turn away from the eye that experiences the greater reduction in light intensity^39^. Whole-field brightness change magnitudes may determine whether animals should swim forward or make a turn^34^. Brightness levels in each eye may also be compared with a dynamically evolving eye-specific luminance set point to determine turn direction^40^. Focusing on purely temporal aspects of the behavior, navigational strategies could also employ the tendency of animals to repeat the same turn direction upon sudden decreases in brightness, thus enabling the return to the original location^37^. However, none of these previous works have systematically validated their models over multiple stimulation configurations, in more complex environments, and across the ontogenetic timeline.

In this study, we use phototaxis in zebrafish transitioning from the larval to the juvenile stage. We find a fundamental shift in behavior as animals mature: larvae spend more time in bright areas, whereas juveniles spend more time in the dark. Using specifically designed visual stimuli, we probe the influence of luminance on swim statistics. Separating temporal and spatial features enables us to disentangle three parallel sensory-motor pathways and build a universal computational model of phototaxis, which allows generating testable predictions in new brightness environments. Our proposed model indicates that animals transition from a spatial contrast comparison to a whole-field luminance averaging strategy. Combining high-throughput tracking experiments with computational modeling in the context of zebrafish phototaxis, our study provides a general framework to describe how behavioral strategies evolve during ontogeny.

## RESULTS

### Inversion of phototactic behavior during zebrafish development

To establish how zebrafish phototaxis changes across two developmental time points, we characterized the behavior of larval (5 days-post-fertilization; dpf) and juvenile zebrafish (26–27 dpf) in different brightness environments. We placed fish in opaque watch glasses, where they could swim freely while their body position and orientation were continuously tracked at high framerates (**Figure 1A**). Our setups allow for the presentation of well-controlled visual stimuli—either locked in closed-loop relative to the fish or in a static open-loop configuration. Carefully designed experiments using such stimuli enable us to build, constrain, and validate behavioral algorithmic models of phototaxis across development (**Figure 1B**). To probe behavioral responses to light, we used four static luminance stimuli with different spatial characteristics (**Figure 1C**): a *half-circles stimulus* with distinct brightness levels on each side (dark vs. bright); a *circular gradient stimulus* that decreased from bright to dark from 0 to 180 degrees and then increased back to bright from 180 to 360 degrees; an *inward gradient stimulus* where brightness increased from the rim towards the center; and an *outward gradient stimulus* where brightness increased from the center to the rim. We tracked individual fish over time (**Figure 1D, Video S1**), allowing us to compute the fraction of time animals spend at different locations across the arena (**Figure 1E**). We also performed experiments in a uniform luminance *control stimulus* (all gray), showing largely homogeneous position distributions for larvae and juveniles (**Figure 1F**, **Figure S1A**). This experiment allowed us to correct for the geometry of the dish by computing values relative to control conditions (**Figure 1G**).

**Figure 1.**
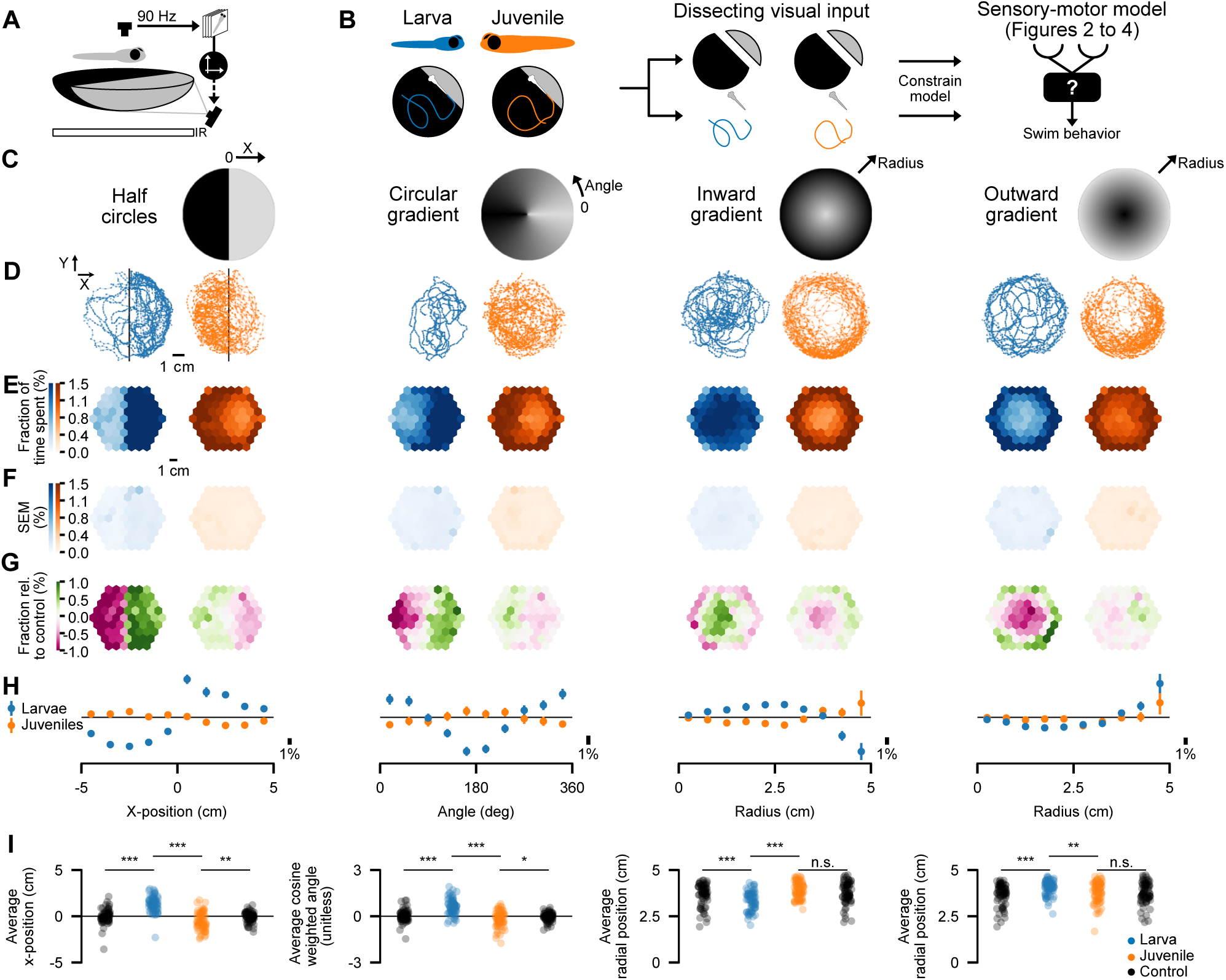
Zebrafish invert phototactic behavior while growing from larval to juvenile stage. (**A**) Experimental setup: The position and orientation of freely swimming fish are tracked in real-time using infrared illumination and a camera. Brightness stimuli are projected from below onto shallow watch glasses (12 cm diameter) using a video projector. (**B**) Our conceptual framework for model-based dissection of ontogenetic switching of visual behaviors. (**C**) Four static test stimuli: *half-circles stimulus* (10 vs. 300 lux), *circular gradient* (300 to 10 lux, and back), *inward gradient* (10 to 300 lux), and *outward gradient* (10 to 600 lux) *stimuli*. (**D**) Example trajectories for a single larva and juvenile (same two animals across stimuli). (**E**) Two-dimensional distribution of the fraction of time spent (%) across arena locations (calculated per fish, then averaged). (**F**) Standard error of the mean (SEM) values for data in (E). (**G**) Same as in (E) but relative to the *control stimulus* (uniform 300 lux across the arena, **Figure S1A**). (**H**) Spatial distributions as a fraction of time (%) spent at arena locations, binned using stimulus-specific metrics (see axis labels in C), relative to the *control stimulus*. Mean ± SEM over fish. (**I**) Stimulus-specific metrics for individual fish. For the *circular gradient stimulus*, angular positions are cosine-weighted. N = 70 larvae (blue) and 73 juveniles (orange). Each fish was presented with all stimuli. Black circles in (H) indicate control conditions (larvae, left; juveniles, right). Annotations indicate statistical significance (p*** < 0.001; p** < 0.01; p* < 0.05; n.s. >= 0.05; Mann-Whitney U-test comparing test stimuli for larvae vs. juveniles as well as the test stimuli vs. control within each age group). See also **Figure S1**.

Across our four tested stimuli, we observed that larvae spent more time in brighter areas (**Figure 1E– G**). Juveniles were more likely to be found in darker regions. For the *outward gradient stimulus*, this effect was weaker. We further computed occupancies along stimulus-specific axes, allowing for individual fish analyses (**Figure 1H** and **1I**), confirming the switch in phototactic behavior from larval to juvenile stage across conditions. The effect sizes between the two age groups were large (Cohen’s d of 2.32 for the *half-circles stimulus*, 3.19 for the *circular gradient stimulus*, 2.57 for the *inward gradient stimulus*, and 0.99 for the *outward gradient stimulus*). Effect sizes compared to the *control stimulus* for larvae were large (Cohen’s d of 2.04 for the *half-circles stimulus*, 2.86 for the *circular gradient stimulus*, 1.16 for the *inward gradient stimulus*, and 1.4 for the *outward gradient stimulus*). Whereas juveniles show large effect sizes for the first two stimuli (Cohen’s d of 0.65 for the *half-circles stimulus*, 0.81 for the *circular gradient stimulus*), the effect size for the *inward gradient stimulus* was medium (Cohen’s d of 0.52) and small for the *outward gradient stimulus* (Cohen’s d of 0.09). The weaker phototactic behavior in the radial stimuli for juveniles could be explained by wall interactions (**Figure S1B**). We speculate that walls may modulate the sensorimotor process in juveniles more strongly than in larvae. As juveniles swim faster than larvae, processing of temporal gradients near the wall may work less well, which could explain our observations. We did not observe body size-dependent effects within each age group, supporting the idea that the ontogeny of the behavior depends on neural rather than bodily development (**Figure S1C**).

Thus, we find a robust behavioral switch in phototaxis behavior across a range of visual stimuli as animals mature from the larval to the juvenile stage. These results hint towards specific adjustments in the behavioral algorithms that animals employ during development. A possible strategy may be based on swimming modulation as a function of whole-field luminance. To probe this idea, we next developed a stimulus that allowed us to measure behavioral responses to varying but spatially uniform brightness levels.

### Modulation of swimming behavior by whole-field brightness

Phototactic behavior may be controlled by static non-spatially dependent whole-field luminance levels (general brightness of the scene), global or localized temporal luminance changes, spatial cues across the eyes, or combinations of these features. The presented static stimuli (**Figure 1C**) revealed a fundamental switch in behavior across development. However, such experiments do not allow for disentangling which of these components are the most relevant ones for the observed effects because spatial and temporal cues vary simultaneously as the fish swims through the arena. To address this problem, we developed a set of well-controlled stimulus conditions to precisely probe temporal and spatial features independently from one another.

We first sought to explore the influence of homogenous, ambient, whole-field luminance levels on behavior. To this end, we developed a virtual phototaxis paradigm with spatially uniform brightness. We picked the brightness level in real-time based on the position of the fish, using the *real circular gradient stimulus* as a lookup function (**Figure 2A, Video S2**). This stimulus mimics the overall time scale of fluctuations when animals move through the arena, while not displaying any spatial contrasts. If animals rely only on spatial luminance cues to guide sensory-motor decision-making, then phototactic behavior should disappear in this configuration, because animals never experience any luminance contrast across the two eyes. On the contrary, if they use an averaging strategy of luminance over the visual field, their phototactic behavior should remain intact. Using the same stimulus-specific metrics as before (**Figure 1G**), we find that larval ability to stay in brighter areas drops to almost chance levels, while juvenile performance improves (**Figure 2B**).

**Figure 2.**
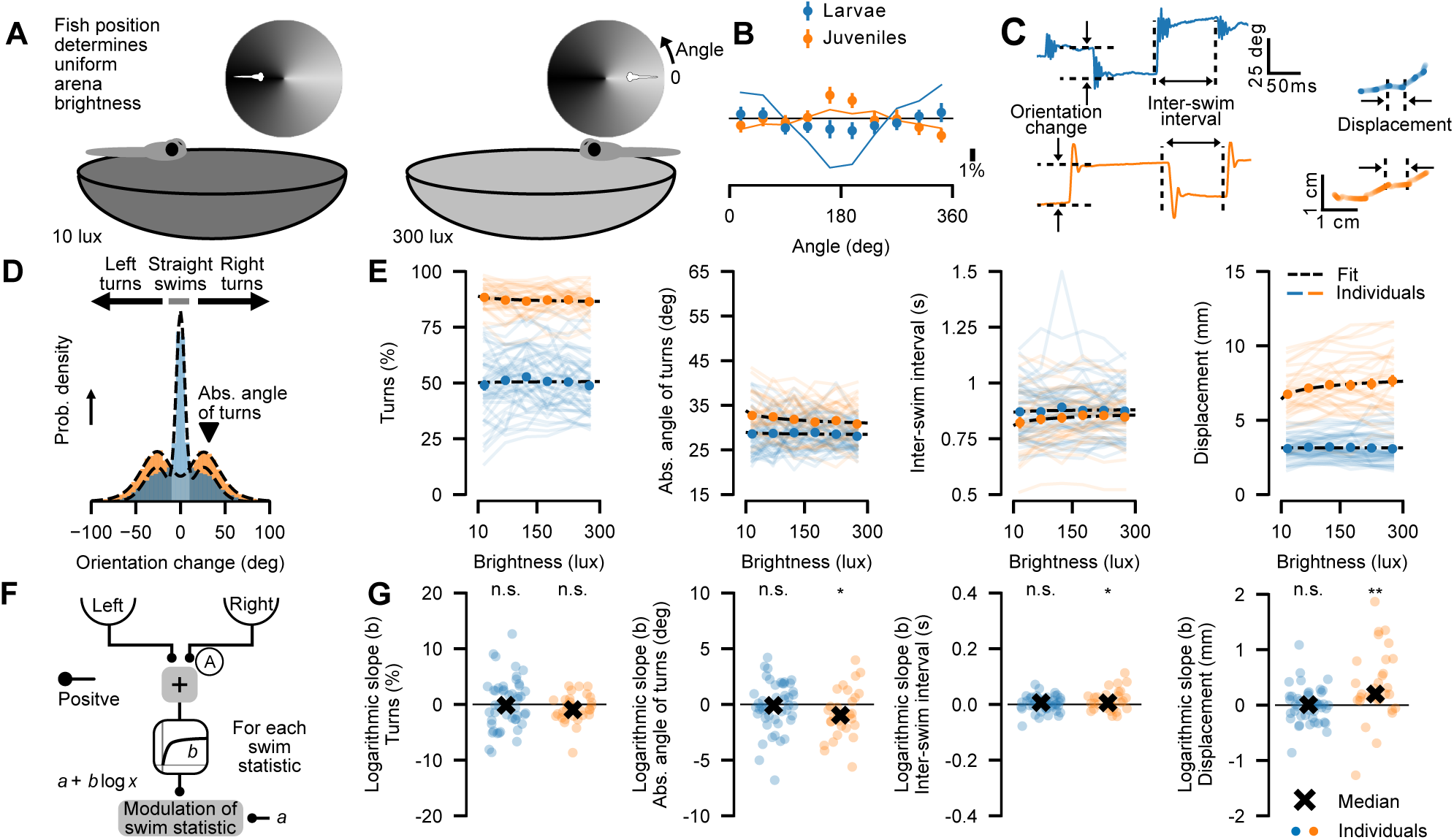
Whole-field brightness modulates swimming statistics in juveniles, but not in larvae. (**A**) Fish are shown spatially uniform brightness levels that are dynamically updated in real-time based on the position of the animal in a *virtual circular gradient stimulus*. (**B**) Same as in Figure 1G, but for the *virtual circular gradient stimulus*. Compared to the spatially distributed *real circular gradient stimulus* (thin lines), luminance navigation performance is largely impaired in larvae, while it improves for juveniles. Mean ± SEM over fish. (**C**) Swims are detected based on sudden changes in heading orientation and quantified by three metrics (orientation change, inter-swim interval, and displacement). (**D**) Normalized distribution of orientation change for larvae and juveniles, and definitions of turns and straight swims. The black arrowhead indicates the absolute angle of turns. Dashed black lines are multimodal distribution fits (Methods). For radius-dependent distributions, see **Figure S2A**. (**E**) Swimming behavior as a function of uniform brightness: Percentage of turns relative to all swims, absolute angle of turns, inter-swim interval, and displacement. Mean ± SEM over fish. Thin colored lines are medians within individual fish, and the black-dashed line is the average over logarithmic fits of individual fish. (**F**) Averaging pathway (A) model (one separate model for each swim metric). This pathway symmetrically averages the luminance over the two eyes and transforms signals into behavioral modulations via a non-linearity. (**G**) Fitted logarithmic slopes for individual fish (circles) of the same data as in E. The black cross is the median over individual slopes. N = 46 larvae (blue) and 30 juveniles (orange). Annotations indicate bootstrapped significance from zero (p** < 0.01; p* < 0.05; n.s. >= 0.05; Methods). See also **Figure S2**.

To further explore the influence of homogenous whole-field luminance on the statistics of swimming, we analyzed behavior from the perspective of individual swims. Both larvae and juveniles exhibit burst-and-glide swimming, allowing us to characterize each event by orientation change, inter-swim interval, and displacement (**Figure 2C**). Larvae and juveniles display symmetric orientation change distributions (**Figure 2D**), as expected for uniform environments. Whereas larvae follow a trimodal distribution, with an emphasis on forward swims, juveniles predominantly exhibit turns, resulting in a bimodal distribution. These distributions were largely independent of radial position in the arena, and thus their shapes cannot be explained by wall-following behavior (**Figure S2A,B**). We classified each swim as a turn when the absolute orientation change exceeded 10 degrees, a threshold corresponding to two standard deviations of the normal peak for larval orientation changes. Otherwise, swims were labeled as straight swims. We used these classifications to compute the percentage of turns relative to all swims and the median absolute angle of all turns.

The extracted swimming metrics allowed us to quantify the effect of uniform brightness levels on behavior (**Figure 2E**) and develop a computational model (**Figure 2F**). This model uses an Averaging pathway (A) in which the surrounding (ambient) luminance is simply averaged and then transformed through a logarithmic nonlinearity into behavioral properties. The averaging first happens within each eye, and then the signals are averaged again. A logarithmic relationship captures the data better than a linear fit (**Figure S2C**), which is in agreement with the Weber-Fechner law^43^. Fitting this model to our experiments allowed us to quantify brightness-dependent modulation across individual fish (**Figure 2G**). For juveniles, we observed that with brightness, the absolute angle of turns decreases, the inter-swim interval increases, and the displacement per swim increases. Overall, these modulations should result in straighter trajectories using fewer but stronger swims to leave brighter areas more efficiently. In contrast, larvae did not significantly adjust any of the measured swim properties based on overall brightness. These results are in line with our observations of animal performance in the *virtual circular gradient stimulus*, where larvae failed to follow the luminance gradient, but juveniles performed well (**Figure 2B**). Thus, our results suggest that larvae do not navigate brightness environments based on a luminance averaging strategy, while juveniles do.

Notably, we observe an enhancement of phototaxis for juveniles in the virtual setting compared to the *real circular gradient stimulus* (**Figures 2B** and **1H**), which hints towards a specific field-of-view for the luminance averaging process. To illustrate this logic, one may assume an extreme case in which juveniles average over large parts of the arena. In such a case, one would obtain similar brightness perceptions independent of their position. If this were true, we would expect uniform distributions of juveniles across the arena for all static stimuli. However, this is not the case (**Figure 1H**). In another extreme, animals may sample only directly under them. In this case, the perceived brightness levels in the virtual and real configurations would be identical, leading to the same performance in the two cases. However, juveniles perform better in the virtual configuration (**Figure 2B**). Therefore, a highly localized averaging strategy is not present either. We conclude that juveniles sample luminance within an intermediate range around their body. We experimentally quantify the spatial extent of this visual range below as part of our agent model design (**Figures 4** and **S4**).

As animals tend to repeat the same turning direction with consecutive swims^44^, we argued that motor history may perhaps also influence phototactic behavior^37^. We found such effects to be weak in our assays for both larvae and juveniles (**Figure S2D** and **S2E**), indicating that motor history-dependent dynamics are likely not the main drivers of phototaxis.

After having explored the influence of homogenous ambient whole-field luminance levels on behavior, we next asked to what degree whole-field luminance temporal changes may modulate swimming. To this end, we tested a stimulus in which the complete arena linearly darkened or brightened in time at different speeds (**Figure S3A**). We chose a broad range of speeds to also cover extreme scenarios. For both larvae and juveniles, we find modulations of swimming behavior only for large whole-field brightness changes (**Figure S3B**). For example, larvae increased their turning angle for strong brightness decrements, reminiscent of a dark-flash response^45^. Juveniles did the same, but for strong brightness increments. Yet, within the intermediate luminance change regimes, modulation of swimming was rather small. We therefore argue that luminance changes are likely less relevant to guide phototaxis under realistic conditions. Based on the observed behavior, we developed a model using an Averaging-Derivative pathway (AD) (**Figure S3C**), in which swim statistics are determined by changes in whole-field brightness levels with different linear transformations for luminance decreases and increases. We fit this model to our experimentally observed swim modulations (**Figure S3B**) to obtain negative and positive slopes for individual fish (**Figures S3E** and **S3F**). A linear relationship captures the data better than a logarithmic fit (**Figure S3D**). Using agent simulations, we will later further quantify the performance of this model (**Figure 4** and **Figure S5G**) by letting the model predict what animals would do under the conditions tested before (**Figure 1**).

In summary, our experimental dissections using the homogeneous whole-field luminance paradigm suggest different strategies for how larvae and juveniles navigate spatial brightness gradients, like the ones tested in **Figure 1**. Juveniles can use the temporal dynamics of ambient luminance cues in a region below them (not requiring any spatial structure). However, larvae do not seem to rely on such cues, suggesting that phototaxis at this early development stage may involve spatial computations across the two eyes. We next developed a range of well-controlled spatio-temporal luminance configurations to explore this possibility.

### Modulation of turns by lateral brightness stimuli

We have shown that larval phototactic behavior is largely absent in our *virtual circular gradient stimulus* (**Figure 2B**) and that behavior is not modulated by homogenous whole-field brightness (**Figure 2E** and **2G**). We thus hypothesized that larvae use spatial contrast cues across the eyes to perform phototaxis in the spatial gradients tested in our open-loop experiments (**Figure 1**). For juveniles, we found that they can perform phototaxis using only whole-field luminance cues (**Figure 2B**). The average brightness around their field of view modulates their swimming statistics (**Figures 2E** and **2G**). However, this result does not rule out the possibility that juveniles also employ spatial information in more complex environments. To explore how these spatial aspects influence directed turns, we implemented a range of visual stimuli in which luminance cues on either lateral side of the fish could be independently controlled. To achieve this, while allowing animals to freely move through the arena, we operated our tracking setup in a real-time closed-loop configuration, such that stimuli are continuously locked to the position and orientation of the fish (**Figure 3A and Video S3**).

**Figure 3.**
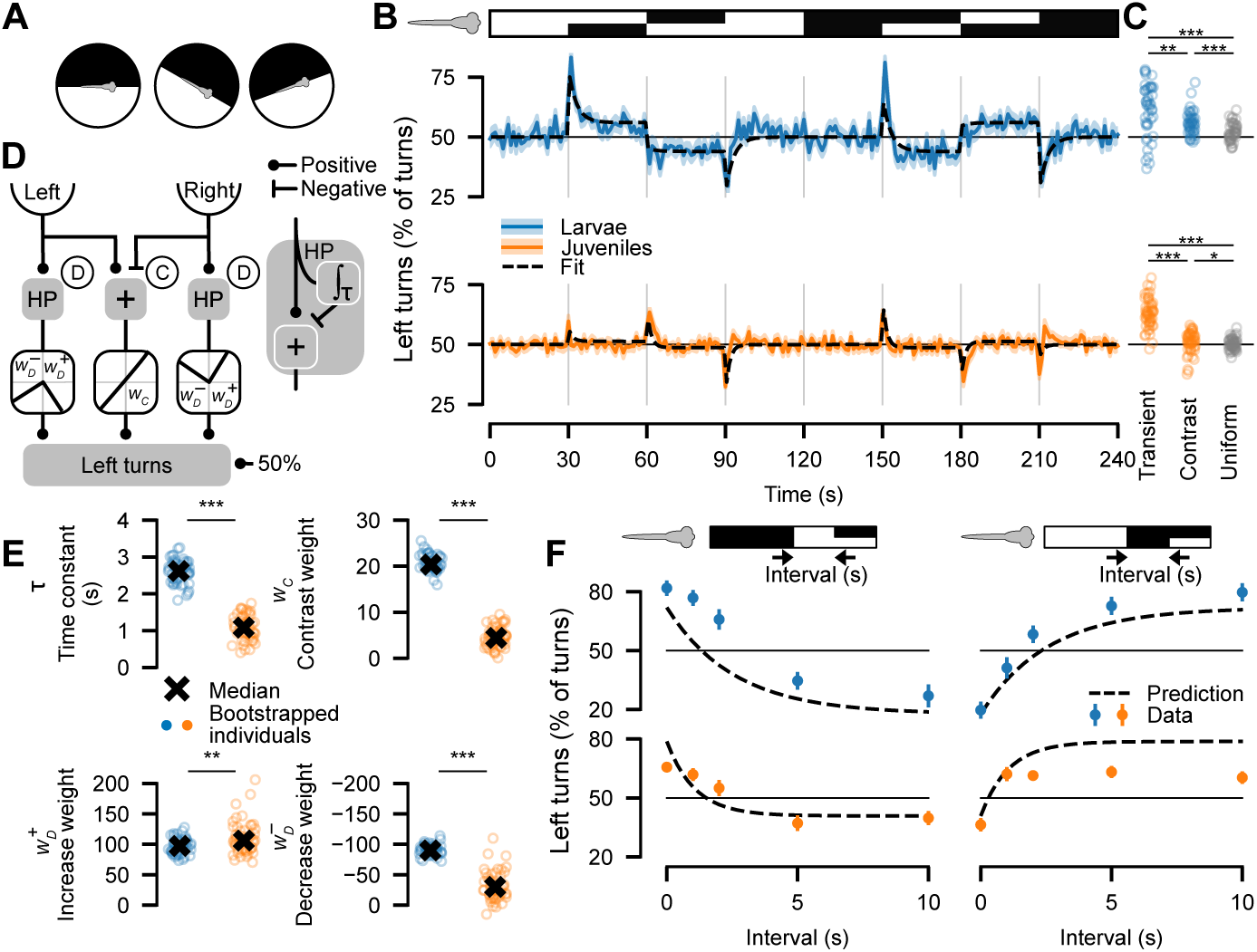
*Lateral brightness stimuli* modulate turn direction. (**A**) *Lateral brightness stimulus*: Visual stimuli are updated based on the position of the fish in real-time, allowing independent control of brightness on the left and right sides of the fish (closed-loop configuration). (**B**) Left turns (%) over time in response to changing lateral brightness levels as depicted in the stimulus bar (top). Black dashed line: mean over individual model fits (see D, E). (**C**) Quantification of data in B for transient, contrast, and uniform phases (Methods). (**D**) Spatial-temporal model: A bias for left turns is determined by a Derivative (D) and Contrast (C) pathway. Increasing and decreasing brightness levels are detected with a high-pass filter. (**E**) Fitted model parameters for individual bootstrapped sets of fish (circles; Methods). The black cross shows the median over all fits. (**F**) Model prediction and experimental validation. Fish are adapted to one uniform brightness, then exposed to a new brightness for a variable interval (left: first 10 lux then 300 lux; right: first 300 lux then 10 lux), and subsequently presented with a dark-bright choice. Black dashed lines indicate model predictions. Quantification is based on left turn peak heights after the transition to the spatial contrast. Experimental data show mean ± SEM over fish. B, E: N = 45 larvae (blue) and 52 juveniles (orange), bootstrapped for E. F: N = 63 larvae and 49 juveniles. Annotations indicate statistical significance (p*** < 0.001; p** < 0.01; p* < 0.05; Mann-Whitney U-test comparing larvae vs juveniles).

Specifically, we designed a *lateral brightness stimulus* consisting of periods of homogeneous brightness, homogeneous darkness, and eye-specific luminance contrast cues locked to the midline axis of the fish (**Figures 3A** and **3B**). We characterized the temporal dynamics of the probability that animals turn left, revealing three major components in the behavioral responses: First, fish tended to transiently turn away from the side that underwent decreasing or increasing changes in brightness (see **Figure 3B** at 30 s and 150 s). Notably, this behavior also occurred when transitioning to a uniform stimulus, even though spatial contrasts are absent during this period (see 90 s and 210 s). These transient responses were present in both larvae and juveniles. Second, larval turning tendencies converged to swimming towards the brighter side when spatial contrast is present (see 50 s, 80 s, 170 s, and 200 s). For juveniles, we found this contrast-dependent behavior to be much weaker and close to chance levels. Third, when luminance contrast abruptly switched sides (see 60 s and 180 s), we found larvae to show a rapid transition between contrast-dependent turning directions while transient turning tendencies were absent. Juveniles, however, transiently swam towards the side that decreased in brightness. We quantified responses at the moment of stimulus transition (= transient), during the spatial contrast steady-state period (= contrast), and during uniform stimulus phases where both hemispheres were either dark or bright (= uniform) (**Figure 3C**). This analysis revealed that both larvae and juveniles respond reliably with directed swims during the transient phases, but that juvenile responses during the steady-state contrast phase were weaker than in larvae.

To quantify latent algorithmic parameters and compare values across larvae and juveniles, we developed a unified model architecture to be fitted to each of the two age groups (**Figure 3D**, Methods). This model is a superposition of two parallel pathways: an eye-specific Derivative pathway (D) and a spatial Contrast pathway (C). This model takes two inputs over time: the brightness on the left and the right side of the fish. For each side, we used a high-pass filter acting like a derivative to detect temporal changes in brightness. We implemented the high-pass filter as a mathematical operation that compares an input signal with its low-pass filtered version (**Figure 3D**, right). We then applied different weight factors, depending on whether brightness increased or decreased. This resulted in a pathway in which luminance changes on the right side induced swims to the left, and the other way around for luminance changes on the left side. The different weight factors can lead to phototactic behavior, favoring increases over decreases or the other way around. We further computed a signed spatial contrast across the eyes by subtracting the brightness on the right side from the brightness on the left side and applied an additional weight factor to the resulting value. Combining these pathways results in a bias in turning direction. The model has only four parameters: the time constant (𝜏) and two asymmetric weights for the Derivative pathway, as well as the weight of the Contrast pathway.

We then fitted this model to our experimental data, allowing us to extract the latent cognitive variables underlying behavior (**Figure 3E**, Methods). Comparison of the resulting values across the larval and juvenile stages then hints toward possible mechanisms of the computational adjustments during development. In the eye-specific Derivative pathway, the time constant was more than twice as high for larvae as for juveniles, suggesting faster temporal processing of visual information in older animals. For both larvae and juveniles, we found the two weight factors for brightness increments and decrements to be of opposing signs. This means that this pathway generally biases turns to go away from localized absolute brightness change for the two age groups. Interestingly, for juveniles, the magnitude of these weight factors was much lower for decrements than for increments, pointing towards asymmetric contrast change computations favoring darkness at this developmental stage (see **Figure 3B**, 60 s and 180 s). For the Contrast pathway, larvae showed a strong positive weight, while the weight for juveniles was almost zero. The cognitive variables differ considerably between larvae and juveniles (Cohen’s d of 4.7 for the time constant, 8 for the contrast weight, 0.62 for the increase weight, and 3.32 for the decrease weight). This result highlights that spatial luminance cues contribute differently in larval and juvenile brightness navigation.

Our model design and its parameters are based on a stimulus that changes on fixed 30-second intervals (**Figure 3B**). We next sought to challenge our model and test its predictive power with new, untested stimuli. To this end, we developed a paradigm in which uniform brightness levels and spatial contrast cues are temporally separated from one another with varying delay intervals (**Figure 3F**). Specifically, we started with a dark uniform environment and then changed the luminance to bright for variable interval lengths. We then switched on the luminance contrast cue, with the bright stimulus on the right side, and probed the initial response during this period. For short intervals, our model predicted that during the luminance contrast phase, turns should be more likely to go towards the dark side, because the model is still adapted to the initial period of darkness. For longer intervals, the model should get increasingly adapted to the bright environment and thus predict more turns towards the bright side. To test these predictions, we performed experiments with fish in the same stimulus configuration. Animal behavior closely matched our modeling results for both larvae and juveniles. As an additional validation step, we repeated analyses on a luminance-inverted variant of the stimulus, where model and experiment were also in close agreement.

We conclude that two more luminance processing pathways are present in zebrafish, in addition to the Averaging pathway discussed in the previous section (**Figure 2**). The Derivative pathway lets larval and juvenile zebrafish transiently swim away from changing brightness levels. The Contrast pathway controls responses to brightness differences across the eyes. Here, when presented with spatial luminance contrasts, larvae swim towards the brighter side, while juveniles remain largely undirected. We have now successfully described behavior in response to well-controlled whole-field (**Figure 2)** and eye-specific spatial-temporal (**Figure 3)** brightness environments. These results enable us to explore whether our fitted models, without further changing their structure and parameters, may predict phototactic behavior when fish dynamically interact with more complex brightness patterns.

### Model simulations of phototactic behavior in complex brightness environments

We have developed algorithmic models for larval and juvenile luminance-based behavioral modulations, whose parameters we fitted to simplified experimental configurations (**Figures 2** and **3**). With these models, we can now test more complex stimuli where spatial and temporal stimulus features are intertwined and where the luminance cues perceived by the animal dynamically depend on its behavior. Moreover, such model simulations allow us to evaluate the relevance of each of the identified pathways for phototactic behavior and how pathway contributions may change across development.

To test our models, we performed agent simulations (**Figure 4A**, **Video S4**), in which model fish navigate stimulus landscapes identical to those presented to real animals (**Figure 1C**). We projected the visual scene onto both eyes of the agent based on its position and orientation in the arena. We fed these time-evolving signals into our models to generate agent movement decisions. As the agents move through the arena, they perceive updated visual inputs. To identify the spatial extent of these projections, we performed experiments in which we varied luminance cues as a function of distance to the animal (**Figure S4A** to **S4C**). We found that swim modulations saturated beyond 2 cm and therefore set the field of view of our agent to a radius of 2 cm.

**Figure 4.**
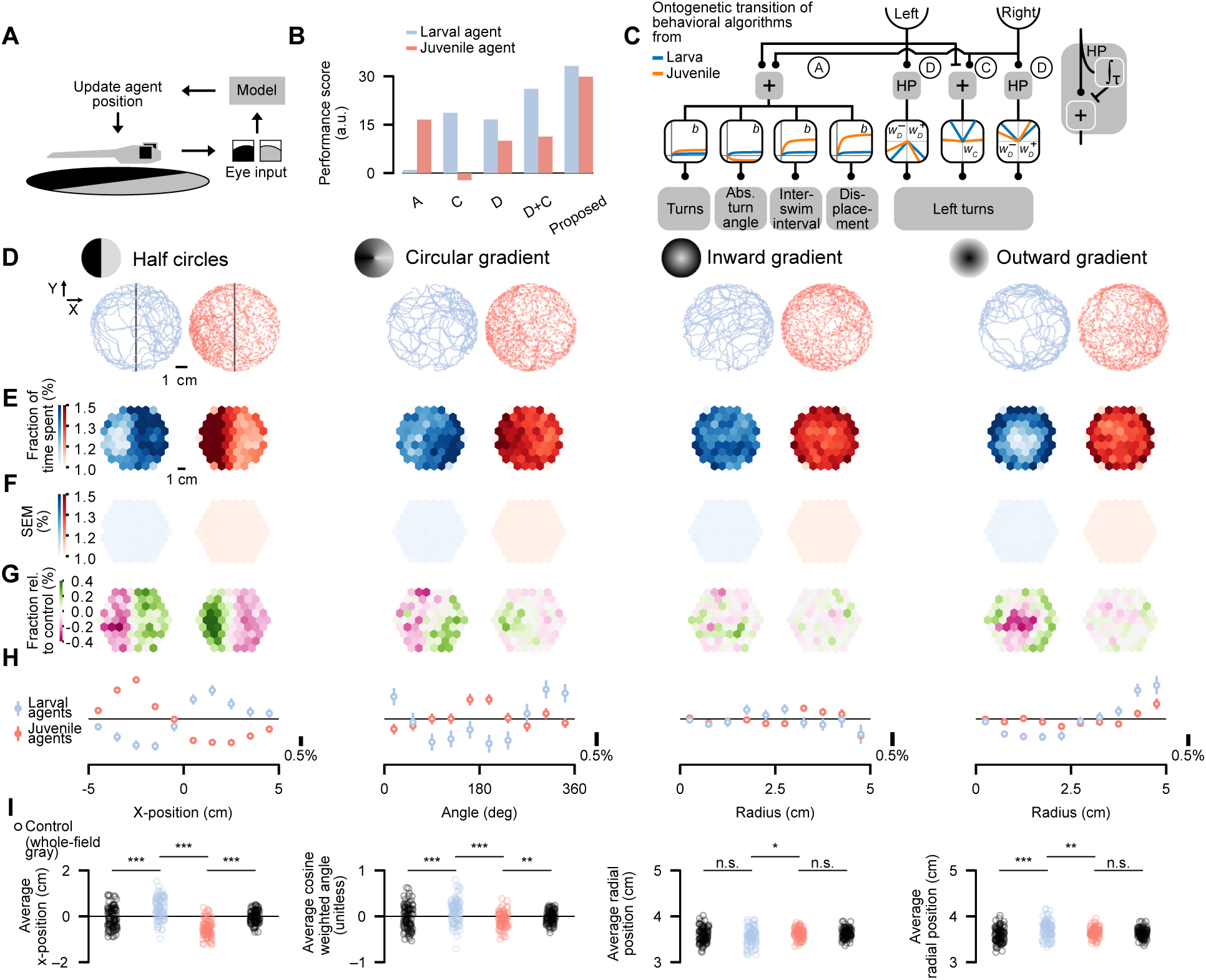
Predicting brightness preferences in complex brightness environments. (**A**) Agents are simulated in virtual arenas of the same dimensions and seeing identical visual stimuli as in Figure 1C. (**B**) Performance scores (Mean-squared error-based), comparing agent spatial distributions to real larval and juvenile data, normalized to blind control agents. Metrics using a Bayesian Information Criterion led to similar interpretations (**Figure S7B**). (**C**) Proposed model architecture: percentage of turns relative to all swims, absolute turn angle, inter-swim interval, and displacement are determined by the Averaging pathway (A, Figure 2F), with turn direction controlled by the Derivative (D) and Contrast (C) pathways (Figure 3D). (**D, E, F, G, H, I**) Same as in Figure 1, but for agent data. N = 96 larva-based agents (blue) and 96 juvenile-based agents (red). Annotations indicate statistical significance (p*** < 0.001; p** < 0.01; p* < 0.05; n.s. >= 0.05; Mann-Whitney U-test comparing test stimuli for larval-based vs. juvenile-based agents as well as the test stimuli vs. control within each age group). See also **Figures S4** to **S7**.

To obtain eye-specific luminance levels, we assume that retinal images are averaged within a certain range around the animal. For juveniles, this assumption is justified by the observation that juveniles show improved phototactic navigation in the *virtual circular gradient stimulus* (**Figure 2B**). To test whether this assumption holds for larvae, we developed another virtual stimulus, the *virtual contrast circular gradient stimulus* (**Figure S4D**). This stimulus is based on the *real circular gradient stimulus* (**Figure 1C**): within a 2 cm radius of the fish, we compute the average brightness on the left and on the right side, and present these two brightness levels as a spatial contrast locked to the midline of the fish. As the larvae swim through the arena, the average luminance to each eye and therefore the perceived contrast will update depending on the orientation and position of the fish. Thus, for the fish, there is never any spatial information within each eye – only spatial contrast across eyes, as well as temporal modulations. As such, this configuration allows us to explicitly test to what extent the luminance input into the Contrast (C) pathway can also be described by local ambient eye-specific averaging. Using this virtual configuration, we found that the spatial distribution of larvae now closely matches the one obtained for the real static circular gradient (compare to **Figure 1H**). This finding, together with our observations that whole-field homogeneous temporal luminance cues are insufficient to guide larvae to the bright locations (**Figure 2B**), thus adds confidence to the design of the proposed Contrast (C) pathway. We corroborate that localized spatially averaged luminance input, followed by an inter-hemispheric comparison, is a driver of phototaxis in zebrafish larvae.

Based on the experimental validations, we ran our agent models as close to the experimental conditions as possible. Specifically, we ran simulations for the same amount of time as for our real fish and generated the same data structure, largely facilitating comparisons between real and virtual animals. Notably, we set the parameters of our agent models solely based on the fitted model results from our separate experiments (**Figures 2** and **3**). This approach allowed us to validate models without fitting them to test stimuli. To compare across different models (**Figure 4B**), we computed a performance score based on the mean squared error between the model and the experimental spatial distributions across the four test stimuli, normalized by the error of a ‘blind’ control model that is fed with constant luminance cues (Methods). A positive performance score indicates that the model simulations show the same qualitative phototaxis tendency as real animals, while a negative value points toward opposing behaviors between the model and experiment.

We first tested the Averaging pathway model (**Figure 2F**). Juveniles show modulations of swimming as a function of whole-field brightness, such as an increased displacement in brighter environments (**Figures 2E** and **2G**). Such a modulation should allow animals to leave brighter areas more rapidly. As larvae did not show such modulations, we expect our larval agent model not to be able to perform the behavior. Our simulations were in agreement with such reasoning (**Figure 4B** and **Figure S5A**).

Next, we tested models composed of the Contrast and Derivative pathways (**Figure 3D**). We first probed each pathway in isolation. The Contrast pathway induces a tendency to swim towards the brighter side for both age groups, with strong effects in larvae but weak effects in juveniles (**Figure 3E**). We expected that such dynamics would be reflected in our agent simulations, which was indeed the case (**Figure S5B**). While our predictions were well aligned with the behavior of real larvae, they were the opposite of what real juveniles do, thus leading to positive and negative model performance scores for larvae and juveniles, respectively. For the Derivative pathway model, our agent simulation qualitatively captured both larval and juvenile behavior (**Figure S5C**), resulting in positive performance scores for both developmental stages. The combination of the Derivative and Contrast pathways (D+C, same model as in **Figure 3D**) further improves the spatial distributions across the four test stimuli for larvae but not for juveniles (**Figure S5D**), reflected by the corresponding positive performance scores.

While these separate pathways in isolation can qualitatively explain individual elements of phototaxis, we require their combination to fully capture the different configurations presented in **Figures 1** to **3**. We therefore propose a model that combines three parallel processing pathways (**Figure 4C**): The Averaging pathway modulates the probability that a swim is a turn, the absolute turn angle, the inter-swim interval, and the displacement per swim. The Derivative and the Contrast pathways then regulate the direction of turns. Not only is this proposed model able to capture the behavioral responses discussed in the previous sections (**Figures 1** to **3**), but it also shows the highest performance score when comparing simulations with the experiment (**Figure 4B**).

To facilitate easy comparison between our proposed model results and experimental data, we plotted simulated data in the same style as displayed before: Our virtual agent simulations mimic the trajectories of real fish (compare **Figures 4D** and **1D**). Looking at the two-dimensional spatial distributions, we observed that simulated larvae spend more time in bright areas, while juveniles go to darker areas (compare **Figures 4E** and **4G**, and **1E** and **1G**). Moreover, our agents largely capture the same stimulus-specific distributions (compare **Figures 4H** and **1H**). We attribute differences in the radial configurations to wall interactions in the experimental data, which we did not explicitly include in our models. Quantifying stimulus-specific metrics for individual fish showed significant differences between larval and juvenile agents (**Figure 4I**), as found in the experiments (**Figure 1I**).

Notably, we obtained these results without tuning any of the model parameters. For juveniles, simulation results and experimental data were in close agreement (**Figure S5E**). In larvae, however, we only obtained qualitative matching of our stimulus-specific metrics (**Figure S5E**). Together, these results indicate that our proposed model likely represents major important pathways in visual processing. To show that our larval simulations would be capable of matching data quantitatively, we next manually tuned one of the parameters in our proposed model. Specifically, we increased the weight of the Contrast pathway in the larval model by a factor of 5. A model with such a configuration mimicked the experimentally observed distributions well (**Figure S5F**).

Finally, we wanted to know if more complex models, capturing additional behavioral features, would perform better than the proposed model. Further analyzing the behavioral response dynamics to the *lateral brightness stimulus* (**Figure 3B**), we found that the other swim statistics, besides the fraction of left turns, were also affected (**Figure S6A**). We therefore designed a five-pathway model that independently modulates each of the five swim statistics (**Figure S6B**): In addition to the Averaging (A), Derivative (D), and Contrast (C) pathways, this extended model also uses an Averaging-Derivative (AD) (**Figure S3C**) and a Derivative-Averaging (DA) pathway. In the latter, we use the eye-specific Derivative pathways and then average signals across eyes. We simultaneously fit our model to the experimental data obtained from both the *lateral brightness stimulus* (**Figure 3B**) and the *virtual circular gradient stimulus* (**Figure 2**), obtaining good matches between model and experiment (**Figures S6A** and **S6C**, **Methods**). We then probed the performance of this extended model in our agent simulations (**Figure S5G**). Although this five-pathway model shows an improvement in the performance score compared to the proposed model, it does not capture the qualitative aspects of the spatial densities better than the proposed model. For completeness, we also tested two models composed only of the Averaging-Derivative pathway (**Figure S3C**) or the Derivative-Averaging pathway, which did not perform better than our proposed model (**Figures S5H** and **S5I**).

In summary, through systematic testing of a variety of different models for phototaxis, we propose a three-pathway configuration regulating swimming behavior. The Averaging pathway controls undirected swimming statistics, while a superposition of a Derivative and a Contrast pathway modulates turn direction tendencies. Even though our proposed model is relatively simple in structure and parameter numbers, it could qualitatively capture behavior across a variety of experimental configurations without parameter tuning. Manual tuning of a single model parameter allowed us to obtain quantitatively matching results. The five-pathway model that modulates each swim statistic separately also performed well, however, with the cost of a largely extended parameter space. Comparing performance score and number of parameters across all tested model arrangements (**Figure S7**), we conclude that our proposed model provides an adequate intermediate level of model complexity and biological realism.

## DISCUSSION

Several studies have explored larval zebrafish phototaxis across a range of stimulus configurations^31–34,37,40^. Yet, there have been no attempts to obtain a comprehensive generalized model that can explain behavior across conditions and over development. Here, we developed a systematic strategy to unravel the algorithmic principles of zebrafish phototaxis, proposing that animals employ three parallel pathways for luminance-based navigation. Our modelling approach allowed us to dissect how computations change as animals become older. It has been reported that juvenile zebrafish tend to swim into darkness, similarly to *Danionella cerebrum* adults^30^. This behavior of juvenile zebrafish is in stark contrast to what larval zebrafish do, which swim into the light, making this transition a suitable example of vertebrate ontogeny.

Although little is known about zebrafish behavior in their natural environment^46^, we hypothesize that eggs are laid in shallow, brighter waters, where the semi-transparent larvae hatch and spend their first days, likely staying hidden from predators or strong currents. As the fish grow and become more visible, they may move to deeper or darker waters for less visual exposure or to start hunting. Another example is the shell-dwelling cichlid, in which the leaving of the nest coincides with an inversion from dark-seeking to light-seeking^47^.

We found that juvenile zebrafish, like larvae, swim in a burst-and-glide pattern. Notably, the two age groups differed in their distribution of turn angles: whereas about half of larval swims are directed straight, juveniles turned more frequently. This increase in turn frequency may reflect a shift in their navigational strategy. With laterally positioned eyes, stronger relative sideways motion could support a better scan of the environment ahead.

While larval swim characteristics showed little sensitivity to uniform brightness levels, juvenile responses to whole-field brightness followed a natural logarithmic relationship, consistent with the Weber– Fechner law^43^. This developmental change may relate to earlier findings that retinal cells in zebrafish keep developing until the juvenile stage^16^. We conclude that whole-field brightness-based behavioral modulations, which can be described via an Averaging pathway, play a major role in the navigational strategy of juvenile zebrafish.

Using closed-loop stimulus delivery methods, previous studies have shown that larvae tend to swim toward brighter regions^33,34^ and away from sides where luminance changes occur^37,40^. We developed this paradigm further to capture both of these behavioral responses within a single experiment. We observed that larval zebrafish behavior is predominantly guided by a Contrast pathway that steers swimming toward the brighter side. Juveniles, however, show very little directional swimming as a function of spatial contrast. Moreover, we find that animals at both developmental stages show a strong tendency to swim away from luminance change, which we describe through a Derivative pathway. This Derivative pathway may be linked to retinal ON-and OFF-pathways, which have been attributed to guide zebrafish larvae along spatial luminance gradients^39^. Yet, as we find that our Derivative pathway induces turns in response to both increments and decrements, it is unlikely to be the only pathway involved in luminance gradient navigation. Instead, it may act as a conservative mechanism to stabilize movement, which becomes particularly notable in high-contrast regions, such as near the midline edge of the *half-circles stimulus*.

Based on our experiments, we propose a three-pathway model for zebrafish phototaxis. Our model includes an Averaging pathway to control the percentage of turns relative to all swims, the absolute turn angle, the inter-swim interval, and the displacement per swim. Derivative and Contrast pathways then bias the turning direction.

Agent-based simulations are powerful tools for developing and validating computational models of behavior and for probing their generalizability^48^. We implemented our proposed model as an agent navigating the same environments as our real fish. Importantly, even though this model was not fitted to the test data, we found that model predictions and experimental results qualitatively match. Comparing this model to a library of alternative model implementations, we further confirm that our proposed model achieves the best predictive performance using only a moderate number of parameters.

Our agent model describes both larval and juvenile behavior using the same algorithmic architecture. We therefore hypothesize that the underlying neuronal connectivity and circuit mechanisms of phototaxis are largely conserved during development. The observed switch from light-seeking to dark-seeking may then be implemented through adjustments in synaptic strengths and potentially rebalancing of pathways. This configuration allows behavior to flexibly adapt to context, internal state, and changing environmental goals, without the need for an energy-consuming restructuring of the brain.

## Limitations of the study

Despite these advances, our study has limitations. In **Figure 3**, we applied bootstrapping over individuals due to the limited number of swim events per fish per time bin, which prevented fitting models to individual fish. Shorter stimulation periods and longer experimental runs may help to mitigate this problem in future experimental designs. Moreover, after manually adjusting the weight of the Contrast pathway, our proposed model could not only qualitatively, but also quantitatively, reproduce larval spatial distributions. Future experiments with a broader range of contrasts may contribute to retrieving a more optimal value of the Contrast pathway weight. Experiments shown in **Figures 1** to **3** were not conducted on the same batches of fish. One could also test the same individuals across all conditions, allowing for single-fish comparisons and individualized model fitting. Furthermore, our tested stimuli are only abstractions of naturalistic visual environments, which may include dynamic motion, depth, and color variation^49,50^. Future work could test whether our findings hold under more ecologically realistic conditions, potentially even in the wild. Our study further assumes that each fish behaves consistently throughout each experiment, which may not be true^51^. Exploring intra-individual variability over time could provide further insights into behavioral flexibility and adaptation.

Although we observe a significant shift in larval and juvenile phototactic behavior, the dark-seeking behavior in juvenile fish appears to be less pronounced than the light-seeking behavior in larvae. One possible explanation is that the juvenile stage examined here represents a transitional developmental phase, in which phototactic strategies are being reorganized but are not fully finalized yet. In addition, experimental constraints may contribute to reduced effect sizes. Juvenile fish swim faster than larvae, which increases the spatial scale over which visual information is sampled and may reduce the effective spatial resolution of the stimuli. It is also possible that phototaxis plays a reduced role at this stage of development, as juveniles may increasingly rely on social or collective cues that are absent in our single-animal assays. Future experiments with broader age ranges, larger arenas, and different luminance configurations will be able to address this issue.

Another limitation of our work is that the identified behavioral algorithms still need to be mapped to their respective neural mechanisms. Zebrafish are well suited for such analyses, on the level of cells and brain-wide circuits, and our results provide the necessary basis for such experiments in the future. Several studies have already hinted at a possible role of the optic tectum in the implementation of the Derivative pathway^39,52^. In larvae, this brain region responds to changes in luminance and plays a causal role during initial turns away from darkness. Long-term phototaxis, over the time span of several minutes, involves intrinsically photoreceptive retinal ganglion cells, the thalamus, and the left habenula^53,54^. Causal manipulations of the habenula influence how likely the fish is to cross a light-dark boundary while leaving swimming speed intact^53^, suggesting a role in the Contrast and/or the Derivative pathway but not in the Averaging pathway. Intrinsically photosensitive neurons in the preoptic area, on the other hand, influence swim statistics but not directional swims^36^, suggesting a role in the Averaging pathway. Future studies probing these brain regions under luminance stimuli, which allow to disentangle the three pathways, could test these predictions. It also remains completely unclear how neural circuits remodel in juveniles to implement the observed behavioral transitions, which should be explored in future work.

In addition, it will be important to quantify behavior along more developmental time points. Such experiments will enable us to better understand if transitions happen gradually or at specific maturation events, for example, with the onset of active feeding. Another promising direction is to study phototactic behavior in animal groups, where social cues may benefit collective decision-making^55^. Here, it will be interesting to combine our experimentally constrained model of phototaxis in juveniles with additional inter-animal attraction rules and probe joint navigational performance. Our modeling framework can be used to make testable predictions for experiments dissecting the structure and function of the nervous system within and across species. For example, it will be interesting to repeat our behavioral analyses in *Danionella cerebrum*, as it has been shown that the response to a light-ON stimulus in these fish is similar to a light-OFF response in zebrafish larvae^56^. This suggests that this species may already be dark-seeking at the larval stage. We have previously used a modeling approach to link specific disease-related mutations to social behavior^21^. Similarly, using recently developed mutant zebrafish with defects in brightness and contrast perception^57^, it will now be possible to test how such manipulations impact our proposed visual processing pathways.

Our work takes advantage of the fact that development and behavior are in an interdependent relationship. We explore two different developmental stages in zebrafish to constrain a universal three-pathway model for phototaxis. This approach allows us to study how decision-making algorithms unfold with development, in organisms that must solve context-specific problems at an earlier stage, while still supporting the development of their behavioral repertoire. In general, our findings provide important new insights into the computational mechanisms that guide vertebrate ontogeny and exemplify the power of model-driven behavioral research.

## RESOURCE AVAILABILITY

### Lead contact

Requests for further information and resources should be directed to and will be fulfilled by the lead contact, Armin Bahl.

### Materials availability

This study did not generate new, unique reagents or materials.

### Data and code availability

● All data associated with this manuscript are publicly available on the persistent repository platform KonDATA with the identifier https://doi.org/10.48606/sb32fkvwj5avwhbr.
● The code for simulations and data analysis is available on the persistent repository platform KonDATA with the identifier https://doi.org/10.48606/m9v18p3c6cb7k77q.
● Any additional information required to reanalyze the data reported in this paper is available from the lead contact upon request.
● The above-mentioned identifiers are already reserved for this paper but will only be available upon final publication. For the review process, please refer to the following link that contains all data and code: https://cloud.uni-konstanz.de/index.php/s/Job9jCGcAe3GjqE.

## ACKNOWLEDGMENTS

We thank the staff of the Animal Facility of the Couzin Lab, and in particular Dominique Leo and Jayme Weglarski, for their generous supply of zebrafish eggs and juvenile fish. In addition, we thank Heike Naumann and Ulrike Bonitz for their vital work in keeping the lab running. We are thankful to the staff support of our central machine shop, especially Ralf Honz, for the invaluable assistance with building our tracking setups. We are grateful for the inspiring and stimulating discussions with Daan Brinks, Iain Couzin, and Osama M. Ahmed. Finally, we would like to thank Daan Brinks, Fabienne Roth, Lukas Pijnacker-Hordijk, Margherita Zaupa, Osama M. Ahmed, and Sophie Aimon for proofreading and providing constructive feedback on the manuscript.

This work was funded by the Emmy Noether Program (BA 5923/1-1), an ERC Starting Grant (101075541 – CollectiveDecisions), and the Deutsche Forschungsgemeinschaft (DFG, German Research Foundation) under Germany’s Excellence Strategy (EXC 2117 – 422037984). In addition, the Zukunftskolleg Konstanz supported A.B. and M.Q.C. The Max Planck Institute for Animal Behavior provided bridge funding for M.Q.C. K.S. was supported by a Boehringer Ingelheim Fonds graduate fellowship.

## AUTHOR CONTRIBUTIONS

Conceptualization: M.Q.C., K.S., P.E.E., and A.B.; Data curations: M.Q.C., K.S., P.E.E., and A.B.; Formal analysis: M.Q.C.; Funding acquisition: A.B.; Investigations: M.Q.C., K.S., P.E.E.,; Methodology: M.Q.C., K.S.; Project administration: M.Q.C.; Resources: M.Q.C., A.B.; Software: M.Q.C., K.S., A.B.; Supervision: A.B.; Validation: M.Q.C., K.S., P.E.E., A.B.; Visualization: M.Q.C.; Writing - original draft: M.Q.C., A.B.; Writing - review & editing: M.Q.C., K.S., P.E.E., A.B.

## DECLARATION OF GENERATIVE AI AND AI-ASSISTED TECHNOLOGIES

During the preparation of this work, M.Q.C. and A.B. used ChatGPT in order to proofread parts of the manuscript and improve language. After using this service, the authors carefully reviewed and edited the content as needed. They take full responsibility for the content of the publication. All illustrations and schematics in the manuscript have been generated by the authors without generative AI, using the vector design program Affinity.

## SUPPLEMENTAL INFORMATION

**Document S1. Figures S1** to **S7**

**Videos S1.** A larval and a juvenile zebrafish in the *half-circles stimulus* with distinct brightness levels on each side (dark vs. bright)

**Video S2:** A larval and a juvenile zebrafish in the *virtual circular gradient stimulus*

**Video S3:** A larval and a juvenile zebrafish in the *lateral brightness stimulus*

**Video S4:** Simulated agent fish in the *half-circles stimulus* with distinct brightness levels on each side (dark vs. bright)

## FIGURE TITLES AND LEGENDS

*Please find all figures with attached titles and legends in the main text*.

## STAR★METHODS

KEY RESOURCES TABLE

KEY RESOURCES TABLE
| REAGENT or RESOURCE | SOURCE | IDENTIFIER |
| --- | --- | --- |
| Deposited data |  |  |
| Data repository | This study | https://doi.org/10.48606/sb32fkvwj5avwhbr |
| Code repository | This study | https://doi.org/10.48606/m9v18p3c6cb7k77q |
| Experimental models: Organisms/strains |  |  |
| Zebrafish: Konstanz wild type strain | Max Planck Institute of Animal Behavior | N/A |
| Software and algorithms |  |  |
| Python 3.12 | https://www.python.org/downloads | RRID: SCR_008394 |
| Panda3D | https://www.panda3d.org | RRID:SCR_021216 |

## EXPERIMENTAL MODEL DETAILS

Wild-type *Danio rerio* (Konstanz strain) larvae, at 5 days post-fertilization (dpf), and juveniles, at 26 and 27 dpf, were used. Larvae were reared in Petri dishes containing E3 solution under a 14/10 hr light/dark cycle at 28°C. Juveniles were housed in ZebTec tanks under a 12/12 hr light-dark cycle and at 28°C. Juveniles were fed dry Zebrafeed by Sparos 3 times a day. In addition, they were fed live high-hufa artemia, according to the following schedule: 5–10 dpf: None; 11–15 dpf: 4x a day 1 drop per feeding; 16–20 dpf: 4x a day 3 drops per feeding; 21–30 dpf: 4x a day 5 drops per feeding. This protocol minimized variation in body size. Care was taken to always include fish from multiple clutches. All experiments were conducted during daytime hours (9 AM to 5 PM). Experiments involving juveniles were approved by the Regierungspräsidium Freiburg under permit number G21/153. To reduce the number of animals used, only zebrafish that were surplus-bred for other projects and could not be used in those projects were included in this study. Sex could not be determined at this age.

## METHOD DETAILS

### Experimental setup

Experiments were conducted in diffusive watch-glass arenas, each filled with 75 mL of E3 solution, resulting in a water volume having a diameter of 12 cm and a maximum depth of 5 mm. The water temperature was stably maintained at 26 °C. Before trials began, fish were given 15 minutes to habituate to the experimental arenas. The visual stimuli were created using Panda3D. Visual stimuli were projected from below using an AAXA P300 Pico Projector. We ensured control over the projected brightness levels by measuring intensities for a fixed interval of projector input values using an LX1330B light meter (Hongkong Thousandshores Ltd., Central Hong Kong). Subsequently, a sigmoid function was fitted to the measured values for each projector and inverted to retrieve the appropriate projector input value for each desired brightness level. Luminance levels varied between 10 lux (dark) and 300 lux (bright) for all stimuli except the *outward gradient stimulus* (**Figures 1C** and **4C**), where brightness increased to 600 lux. For the circular and radial gradients, the brightness scales linearly between this brightness range along the azimuth and radius, respectively. The stimulus position was calibrated to match the camera position, and a computational warping grid was applied to allow orthogonal projection onto the curved watch-glass arenas. Stimuli were shown in random order, and the same fish were exposed to stimulus-corresponding controls. Stimuli were shown either as static or locked to the fish’s position in a closed-loop setup configuration (full closed-loop delay: 60 ms). All stimuli were also always shown in their mirrored version to control for potential asymmetries in behavior and arena shape: around the arena’s y-axis for static stimuli (**Figures 1**, **2**, and **4**) or around the fish’s longitudinal body axis for fish-locked stimuli (**Figure 3**).

Infrared illumination was provided by an LED array (940 nm wavelength), and behavioral recordings were made using an IR-sensitive camera (Basler acA2040-90um-NIR) equipped with an adjustable macro lens (Navitar ZOOM 7000) and an IR longpass filter (Linghuang Zomei IR 850 nm 52 mm). The camera operated at 90 Hz. To increase experimental throughput, we used two computers, each connected to eight cameras and four projectors, allowing for independent stimulation and closed-loop tracking of 16 fish simultaneously. Fish position and orientation were continuously measured using background subtraction-based tracking, implemented in custom Python software. To compute body size (**Figure S1C**), we implemented a real-time posture-based tracking to detect the midline. Experiments, simulations, and data analysis were performed using custom-written Python code (Python 3.11).

### Agent simulations

Each simulation ran at 60 Hz and included 96 model agents. The simulations were run in parallel at the agent level, in which each agent was assigned a unique random number generator seed. The simulated arena matched the real experimental arena (6 cm radius). Simulated agents received identical visual stimuli as the real fish, with the same trial structure, trial duration, and number of trials. Each agent had a left and right eye, each with a 2 cm half-disk field of view, where the half-circle region on the side of each eye was considered for visual input (spatially averaged, 𝐵_𝑟𝑖gℎ𝑡_ and 𝐵_𝑙𝑒𝑓𝑡_). Pixels outside the dish but within the field of view were assigned not-a-number values and ignored in further processing. The stimulus was rendered on a grid of 256 by 256 pixels. At the start of each trial, each agent was assigned a random orientation and position within the arena. The swim events of agents were determined based on the brightness input to the corresponding models. All models had parameters obtained from real fish data (**Figures 2** and **3**, and **S3**), fitted for each age group and swim statistical property.

The Averaging pathway (A) models (**Figure 2D**) took the average brightness (𝐵) across the two eyes

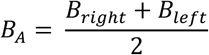

and the contribution to the swim characteristics by evaluating the natural logarithm

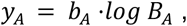

where 𝑏_𝐴_ is the logarithmic slope.

The Contrast (C) pathway models (**Figure 3D**) compute the contrast across the eyes following

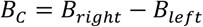

and its contribution to swim characteristics was obtained by evaluating the linear function

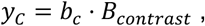

where 𝑏_𝑐_ is a linear transformation.

The Derivative (D, **Figure 3D**), Averaging-Derivative (AD, **Figure S3C**), and Derivative-Averaging pathway (DA, **Figure S5I**) involve a high-pass filter to detect luminance changes. This filter is implemented via a low-pass filter that is slowly updated to match the new brightness value. The low-pass filtered value is updated at each timestep 𝑛 following

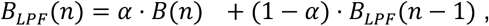

where the low-pass filter parameter 𝛼 is based on the time constant 𝜏 and time since the last processed frame 𝛥𝑡:

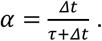

Subsequently, the high-pass filter value is obtained by computing the difference between the current perceived brightness 𝐵(𝑛) and the low-pass filtered 𝐵_𝐿𝑃𝐹_(𝑛) by

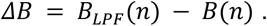

For these pathways, the resulting brightness change value is subsequently multiplied by sign-dependent weights and linearly transformed to obtain the pathway contributions to the modulation of swim characteristics according to

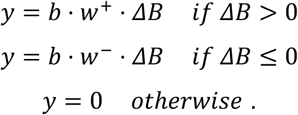

Where scaling factors 𝑏, and weights 𝑤^+^, and 𝑤^−^ were different for each of the D, AD, and DA pathways.

To obtain the final swimming characterizations for all five pathways, the contributions of each pathway are then summed, and an offset 𝑎 is added. For the extended five-pathway model, this results in

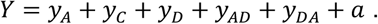

In the proposed model, the percentage of turns relative to all swims, turn angle, inter-swim interval, and displacement are determined by the Averaging pathway only, as given by

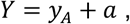

and the fraction of left turns by a superposition of the Derivative (D) and Contrast (C) pathways is given by

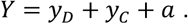

Upon reaching the arena wall, agents maintained their general swimming direction but were assigned a new heading angle relative to the wall, drawn from a normal distribution centered around 115 degrees and with a 10-degree standard deviation. The angle drawing process was repeated until the agents moved away from the wall again.

## QUANTIFICATION AND STATISTICAL ANALYSIS

### Edge interactions, pre-screening, and data exclusion

To exclude edge effects, only data from fish located at least 1 cm away from the edge were included in the analysis. As part of the habituation to the experimental arena, larvae underwent a 10-minute pre-screening period to exclude unhealthy individuals. Larvae that did not adhere to the following behavioral criteria were excluded: Leftward and rightward gratings: at least 50% of turns aligned with the grating direction; Homogeneous gray background: maximum of 20% of turns in a single direction. For juveniles, the arena was left gray during the habituation phase without performing a pre-screening protocol. To exclude potential tracking errors during the pre-screening and the experiment, swim events were discarded if they exceeded predefined thresholds. Generally, less than 1% of all data was discarded. Larvae: maximum contour area = 12,000 px, maximum average speed per swim = 1.5 cm/s. Juveniles: max. contour area = 16,000 px, maximum average speed per swim = 3 cm/s. All fish: maximum inter-swim interval = 10 s, maximum absolute orientation change = 150°. A complete trial was discarded if more than 5% of swim events in that trial exceeded any of the above thresholds.

### Swim event detection

Fish position and orientation were extracted after background subtraction, with the background being updated every 120 s. Swim events were identified based on windowed variance thresholds applied to movement data, using a variance window of 0.050 s, a minimum duration above the start threshold of 0.020 s, a minimum duration below the end threshold of 0.050 s, and a maximal event length of 4 s. The start and end variance thresholds differed between larvae (start threshold = 5, end threshold = 1.5) and juveniles (start threshold = 7.5, end threshold = 5).

#### Figures 1 and 4

The tracking data described the x and y positions for every individual, stimulus, and trial, with each row denoting a frame. Spatial binning was applied using 10 bins. Bins were defined for x-position, radius, and azimuthal angle, with values ranging from –5 to 5 cm for x, 0 to 5 cm for radius, and 0 to 360 degrees for the angle. The spatial distributions were first averaged within individuals before calculating the mean and standard error of the mean over each population. The shown probabilities (**Figures 1, 2B,** and **4**) are normalized by the values of the *control stimulus* (**Figure S1A**). The average cosine-weighted angle is a dimensionless metric used to capture the average position of individuals for all tested *circular gradient stimuli*. This was computed by multiplying the number of swims within each azimuthal bin 𝑁(𝛼_𝑖_) with the cosine-weighted value 𝑐𝑜𝑠 (𝛼_𝑖_) for that bin and subsequently summing over all bins

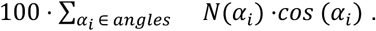

Simulations produced the same format as the experimental data, allowing us to perform the exact same analysis for both figures. The agents’ performance score indicates how well an agent captures the spatial distribution of the real fish. We calculate the average mean-squared error for each of the four (= 𝑁_𝑠𝑡𝑖𝑚_) test stimuli in their stimulus-specific spatial bins, comparing the spatial distributions of simulated agents (𝑝_𝑎g𝑒𝑛𝑡_) to real fish (reference, 𝑝_𝑟𝑒𝑓_)

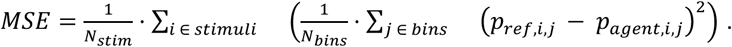

Subsequently, the performance was calculated as the percentage improvement relative to a blind control model (perceiving constant luminance levels),

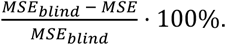

#### Figure 2

The swim-event characteristics (orientation change, inter-swim interval, and displacement) are summarized as median values over trials, within individual fish and stimuli, as the underlying distributions are not necessarily symmetric. Orientation change can be best described by a mixture of a Normal distribution for forward swims and two Maxwell-Boltzmann distributions for left and right turns. The proportions of left and right turns can be captured using weighting factors in the mixture model. Displacement and inter-swim intervals both follow a Gamma distribution, which is known to be well-suited for representing temporal and spatial intervals.

In the virtual circular gradient paradigm, the arena brightness was set based on the azimuthal position of the fish. Swims were grouped into brightness bins, based on the brightness at the start of the swim, and medians were computed within each brightness bin for individual fish. A logarithmic function was then fitted to the brightness data (see below), allowing for an estimation of the relationship between brightness and swim characteristics. To assess whether the fitted slope parameters differed significantly from zero, we performed bootstrapping over fish with 10,000 iterations to compute two-tailed p-values and 95% confidence intervals. This approach provided robust statistical inference for evaluating the modulation of swim parameters by brightness. The Averaging pathway (A) model (**Figure 2D**) is described above in the subsection called ‘Agent simulations’.

#### Figure 3

For the quantification in **Figure 3C**, we divided data points for individual fish into three phases. The transient phase included data points in the first temporal bin after the appearance or disappearance of spatial contrast. The spatial contrast and uniform phase both include the center 10 seconds of the stimulus blocks, where the stimulus shows a spatial contrast or uniform brightness.

To ensure sufficient data points per temporal bin for model fitting (**Figure 3E**), we generated bootstrapped sets of fish. For each age group, the total number of bootstrapped fish matched the number of original individuals. Each bootstrapped fish was constructed by randomly sampling, with replacement, two-thirds of the available real fish. For time-series analysis, data were resampled by calculating the median value within 1-second time windows for each individual. The Contrast (C) and Derivative (D) pathways (**Figure 3D**) are described above in the ‘Agent simulations’ subsection. Models were fitted to individual bootstrapped fish (see below), and mean model parameters were retrieved by averaging over parameters of the individual fits.

To detect peaks in the percentage of left turns (**Figure 3F**), values before stimulus onset were excluded, and the peak was determined by maximizing the absolute deviation from the 50% chance level.

All statistical tests are specified in the figure legends. To compare distributions against zero, we used bootstrapping to compute a two-tailed p-value. Unless specified otherwise, resampling was performed 10,000 times, each time selecting 2/3 of the individuals to create a bootstrapped dataset matching the original sample size. For comparisons between distributions, we applied the Mann-Whitney U-test.

Model parameters were obtained by fitting the analytic model equations to experimental data using the optimization and root-finding functions of the Python package SciPy. Fitting to separate experimental datasets–*virtual circular gradient stimulus* (**Figure 2**), *lateral brightness stimulus* (**Figure 3**), and *whole-field luminance changes stimulus* (**Figure S3**)–was performed using the non-linear least squares fit function *curve_fit*. The extended five-pathway model was fitted to the combined experimental data from the *lateral brightness stimulus* and *virtual circular gradient stimulus* by minimizing the mean-squared error to both datasets using the *minimize* function.

## SUPPLEMENTAL INFORMATION

**Document S1**

**Figures S1** to **S7**

**Figure S1.**
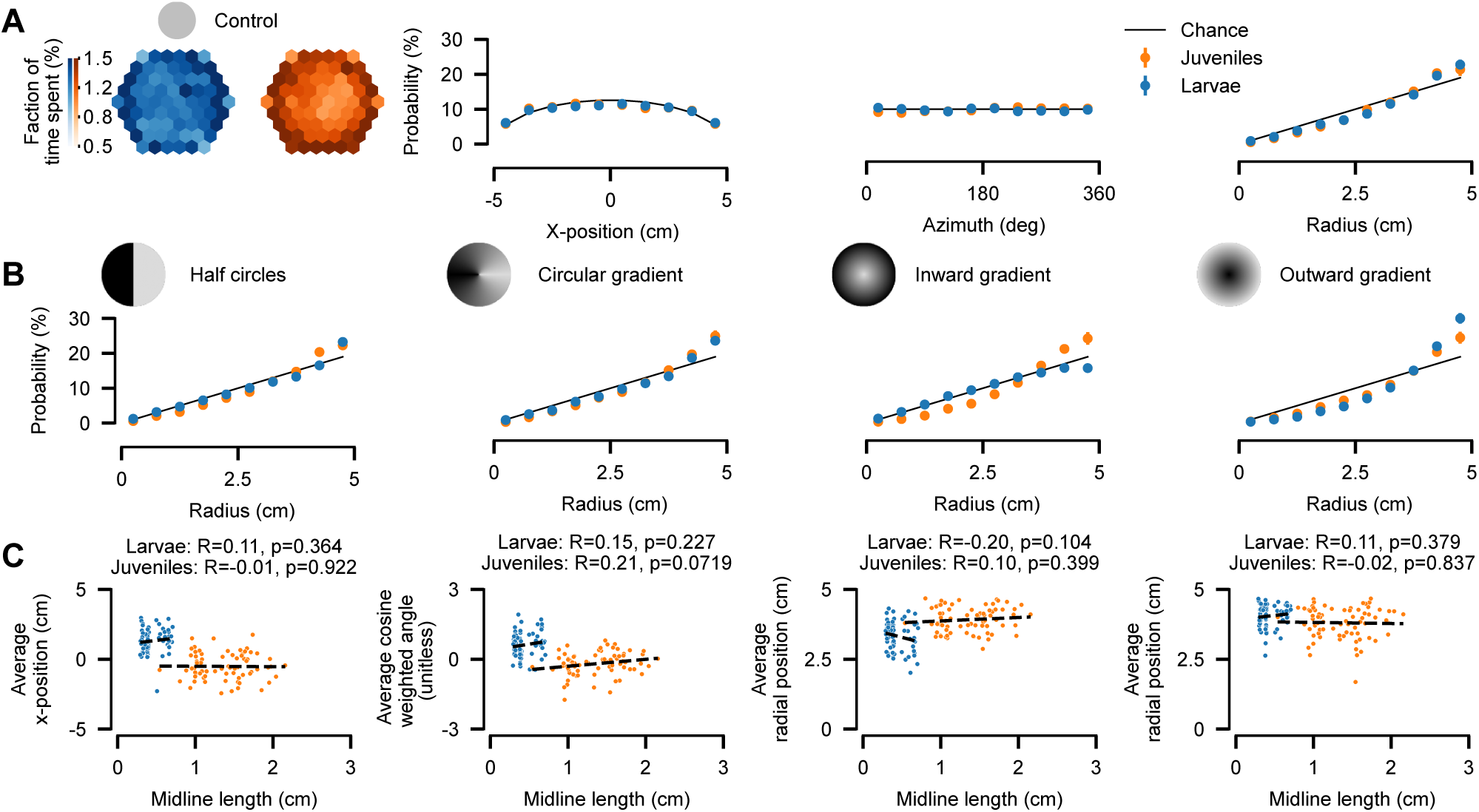
Controls for static luminance experiments. (A) Larvae and juveniles are distributed homogeneously in the *control stimulus* (uniform gray, 300 lux, across the arena). Fraction of time (%) spent at arena locations, binned using the same metrics as in Figure 1G, but not normalized to the *control stimulus*. Mean ± SEM over fish. (B) Arena occupancy as a function of the radial distance to the center for all stimulus conditions. Wall interactions add a small additional attraction towards the rim for the *half-circles stimulus* and *circular gradient stimulus*. For the *inward and outward gradient stimuli*, behavior is a superposition of wall interaction and gradient navigation. (C) The relation between body size (as measured by the length of the midline) and phototactic behavior for individual fish. Body size within each age group does not influence the position of the fish in the arena. N = 70 larvae (blue) and 73 juveniles (orange), thin black lines indicate mathematical chance levels, and black dashed lines indicate linear fits to age populations. Related to Figure 1.

**Figure S2.**
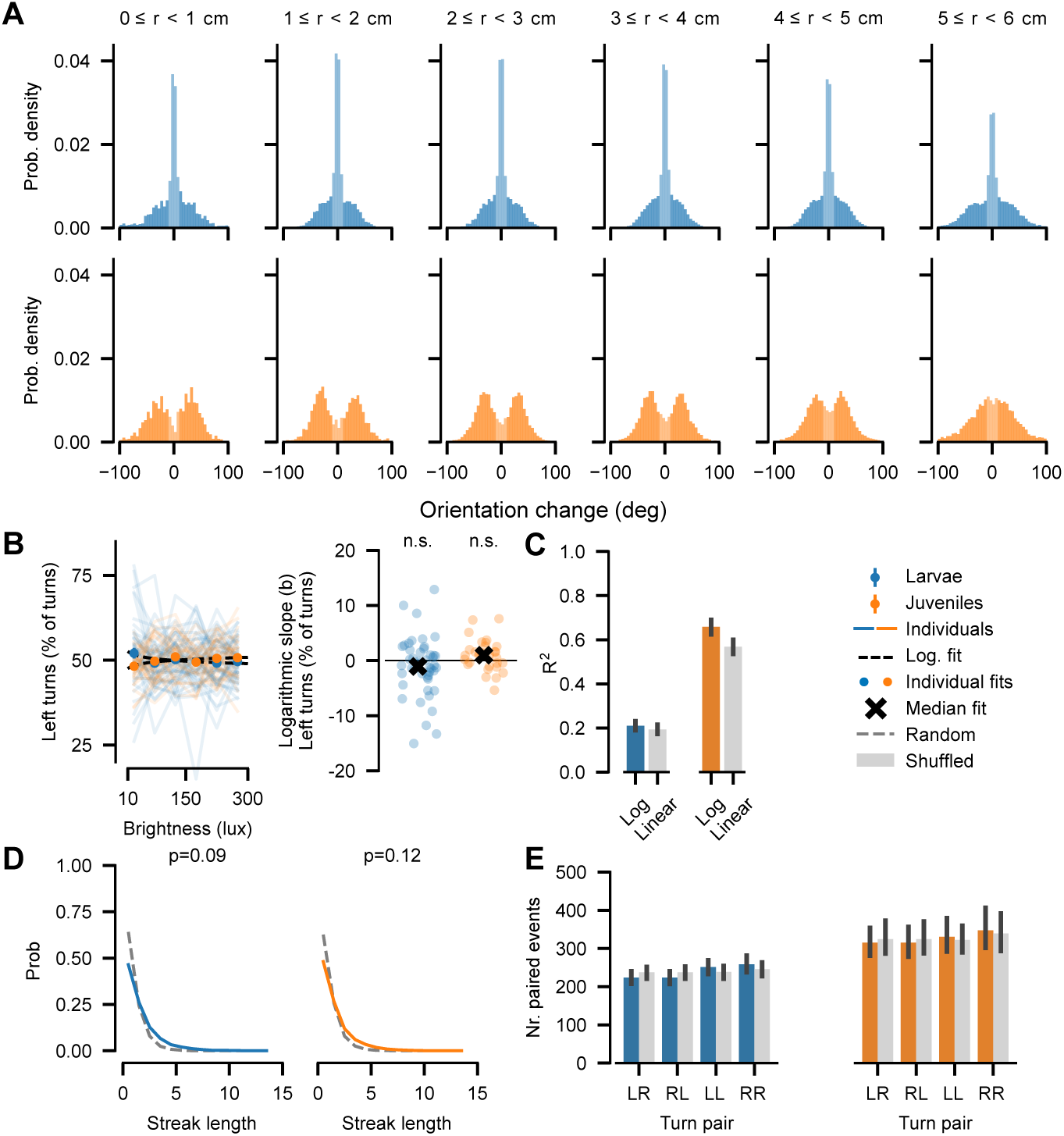
Detailed quantification of behavior in the *virtual circular gradient stimulus*. (A) Area-normalized distributions of orientation changes at different radial bins in the arena. (B) Fraction of left-directed turns as a function of whole-field brightness. Annotations indicate bootstrapped significance from zero (n.s. >= 0.05). (C) A logarithmic relationship of data in Figure 2E captures the modulation of behavior by whole-field brightness for juveniles better (higher R-squared) than a linear relationship. As larval behavior is not strongly modulated by homogenous whole-field luminance, both functions lead to the same fitting quality. (D) Following the analysis performed previously^44^, we only observe a weak, not significant effect in streak length probabilities when compared to randomly generated turns. P-values indicate statistical significance (Mann-Whitney U-test comparing randomly generated turns to real data). (E) Turn pair counting for individual fish does not show a significant difference from shuffled data either. N = 46 larvae (blue) and 30 juveniles (orange). Black vertical bars indicate 95% bootstrapped confidence intervals. Related to Figure 2.

**Figure S3.**
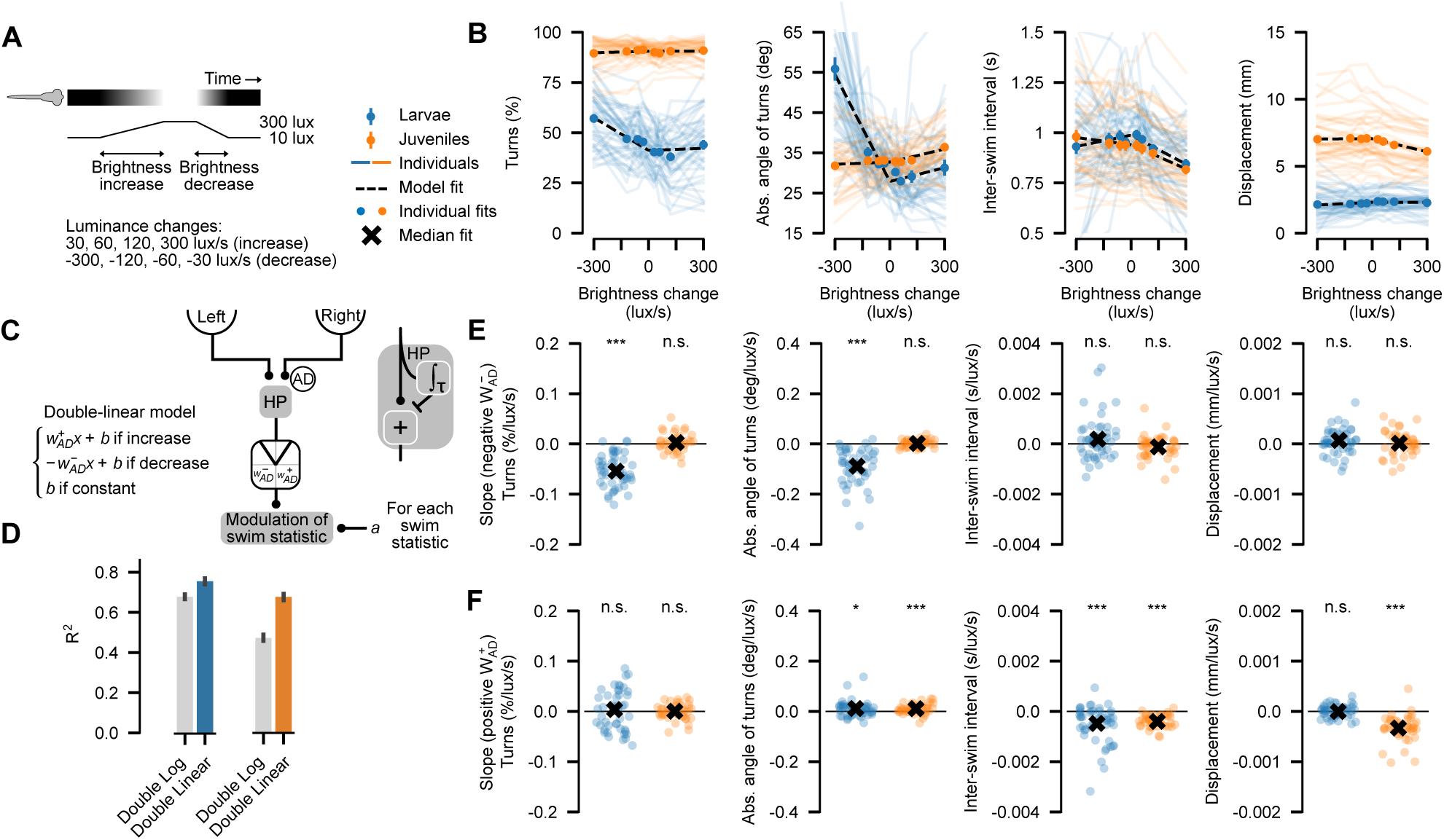
Whole-field luminance changes. (A) We present whole-field luminance changes by linearly increasing or decreasing the arena brightness with either 30, 60, 120, or 300 lux/s. (B) Swimming behavior as a function of temporally changing whole-field brightness levels: Percentage of turns relative to all swims, absolute angle of turns, inter-swim interval, and displacement. Mean ± SEM over fish. Thin colored lines are median values within individual fish, and the black-dashed line is the average over the fits of individual fish. (C) Averaging-Derivative pathway (AD) model. (D) A double-linear relationship (larvae, blue; juveniles, orange) captures the modulation of behavior by whole-field brightness better than a double logarithmic relationship (gray). (E, F) Fitted slopes for individual fish (circles) of the same data as in B. N = 54 larvae (blue) and 43 juveniles (orange). The black cross is the mean over individual slopes. Annotations indicate bootstrapped significance from zero (p*** < 0.005; p* < 0.05; n.s. >= 0.05; Methods). Related to Figures 2 and **3**.

**Figure S4.**
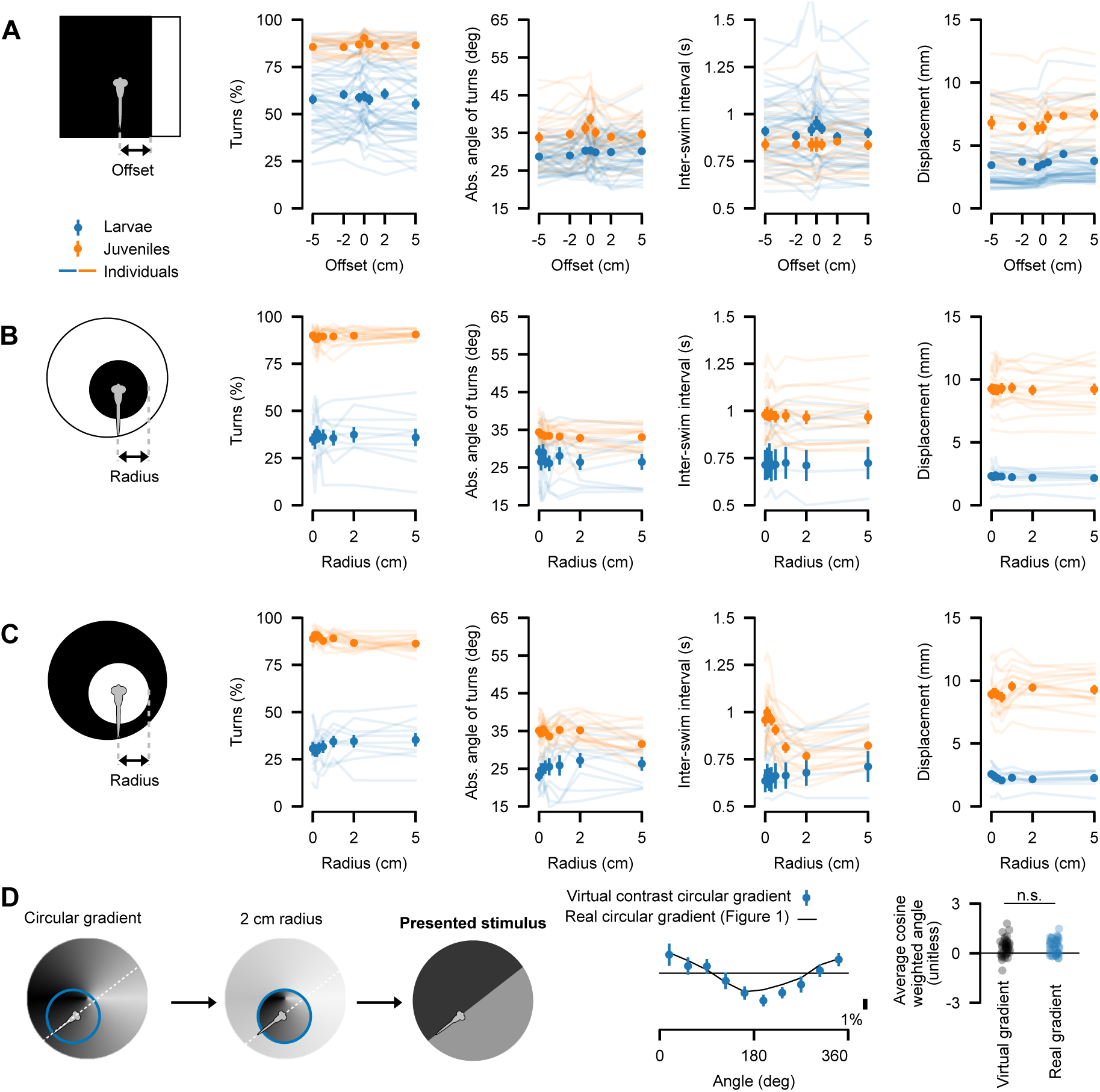
Field of view measurements reveal the spatial extent of luminance calculations. To gain an impression of the field of view of larval and juvenile zebrafish for luminance calculations, we quantified swim characteristics for three different fish-locked stimuli. (A) A stimulus where the lateral offset position of the contrast edge is varied relative to the midline of the fish. (B) A dark disk centered at the position of the fish in a bright environment. (C) A bright disk in a dark environment. For each configuration, we quantified how behavior is modulated by the offset or radius of the stimulus. (D) *Virtual contrast circular gradient stimulus*: Fish are shown a spatial contrast locked to the orientation and position of the fish (right). The brightness levels for the left and right sides are determined by averaging brightness values within a 2 cm radius on either lateral side (middle) of the fish coordinates with a lookup table using the *real circular gradient stimulus* (left, and Figure 1C). Compared to the *real circular gradient stimulus* (thin black line), luminance navigation performance is not significantly different. Mean ± SEM over individual fish, thin colored lines are medians within individual fish. A: N = 47 larvae (blue) and 20 juveniles (orange), B and C: N = 10 larvae and 16 juveniles, D: N = 22 larvae. Related to Figure 4.

**Figure S5.**
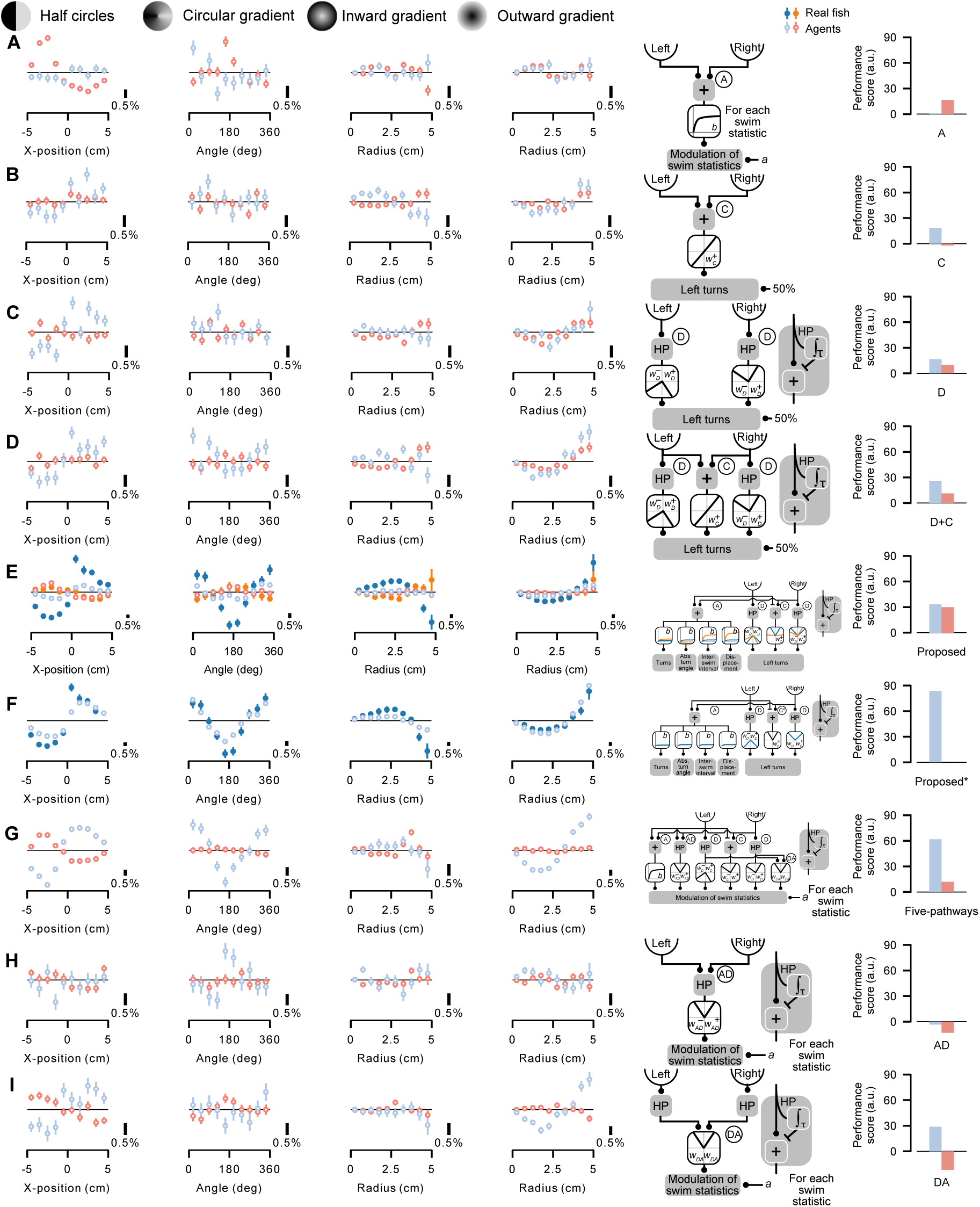
Detailed model predictions for all agent model variants. (A–I) First four columns: Fraction of time (%) spent at arena locations, binned using stimulus-specific metrics (see axis labels in Figure 1C), relative to the *control stimulus* (all gray). Mean ± SEM over agents. Fifth column: model illustration. Sixth column: Performance score. A to I: Averaging pathway model (A), Contrast pathway model (C), Derivative pathway model (D), Derivative and Contrast pathway model (D+C), the proposed model (Proposed), the proposed model (Proposed*) with manually tuned Contrast weight (larval agent only), Extended model (Five-pathways model), Averaging-Derivative pathway model (AD), and Derivative-Averaging pathway model (DA). N = 96 agents for each model simulation and age group, larval-agents (hollow light blue), juvenile-agents (hollow red), 70 real larvae (solid blue), and 72 real juveniles (solid orange). Related to Figure 4.

**Figure S6.**
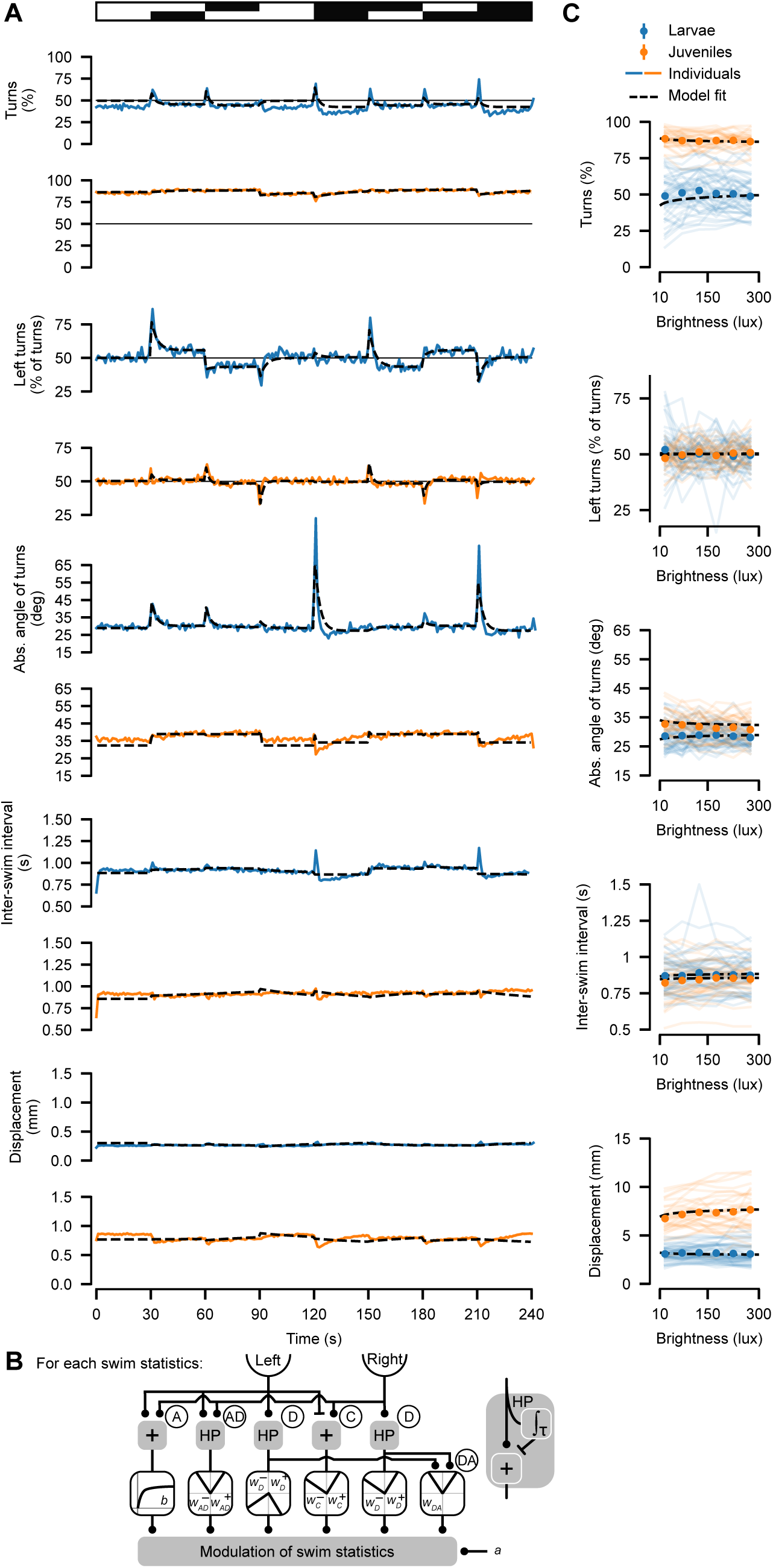
The extended Five-pathways model for the *lateral brightness* stimulus. (A) Swimming behavior in response to the *lateral brightness stimulus* (Figure 3B): Percentage of turns relative to all swims of all swims, fraction of left turns of all turns, absolute angle of turns, inter-swim interval, and displacement over time. (B) Extended model including all pathways: Averaging (A), Averaging-Derivative (AD), Derivative (D), Contrast (C), and Derivative-Averaging (DA) pathway. (C) Swimming behavior as a function of uniform brightness level (Figure 2A). A: N = 45 larvae (blue) and 52 juveniles (orange), C: N = 46 larvae and 30 juveniles. Thin, solid, colored lines are individual fish. Black dashed line: mean over individual model fits. Related to Figures 3 and **4**.

**Figure S7.**
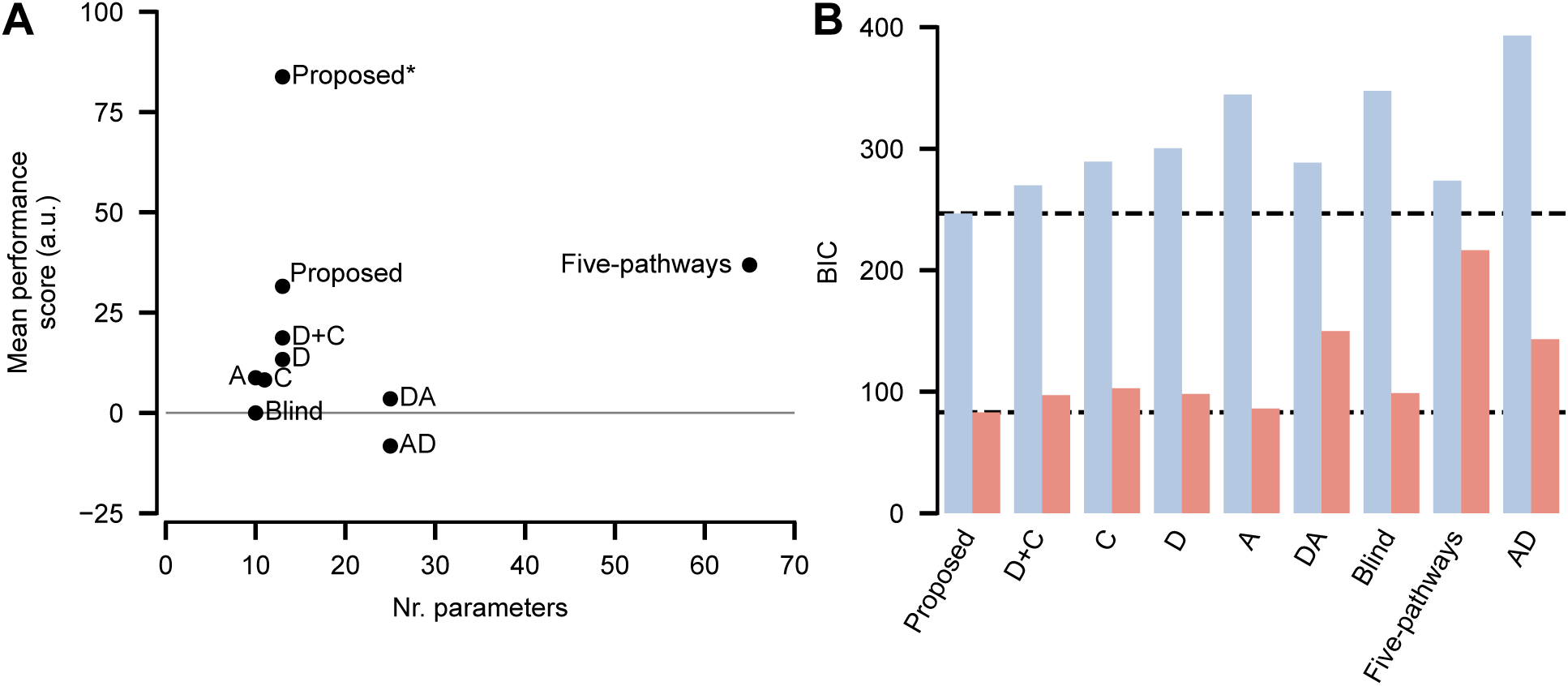
Simulated model performance across all tested agent configurations. (A) Mean performance scores over larval and juvenile-based models vs model parameters. Blind: Constant luminance input, C: Contrast pathway, D: Derivative pathway, D+C: Derivative and Contrast pathway, Proposed: proposed model, Proposed*: proposed model with manually tuned Contrast weight (larval agent only), DA: Derivative-Averaging pathway, AD: Averaging-Derivative pathway, Five-pathways: extended model. (B) Bayesian Information Criterion (BIC) to compare model performance given their number of parameters. For both larvae (blue) and juveniles (orange), the BIC is smallest (= best) for our proposed model. Related to Figure 4.

